# Abnormal bioelectricity during unperturbed cancer growth. Part-I: electrical biopotentials, surface charge density, volumetric charge density, phase-field and Turing approach

**DOI:** 10.64898/2025.12.19.695347

**Authors:** J. Roberto Romero-Arias, Luis Enrique Bergues Cabrales, José Alejandro Heredia Kindelán, Daniel Romero Rosales, Rafael A. Barrio, Juan Bory Reyes, Juan Ignacio Montijano

## Abstract

Understanding the abnormal cancer bioelectricity and its close relationship to growth is a challenge for researchers and oncologists. The objective of this study is to simulate the mechanical-electrical coevolution of solid cancer using an energetic approach, which involves the local and global changes of bioelectricity in the entire unperturbed tumor, aspects that may be correlated spatially and electrically with the growth, progression, nucleation, angiogenesis and metastasis of a unperturbed solid malignant tumor. For this, spherical harmonics, phase-field, and the Turing approach were taken into account in the model. The results showed that electrical bioptentials and volumentric charge density throughout the interior of the unperturbed tumor, as well as surface charge density at tumor-surrounding healthy tissue boundary during its growth depended on mass aggregation, time and type of spherical harmonics. Regions with both positive and negative charge densities throughout the interior of the unperturbed tumor volume were observed. We concluded that positive and negative values may be responsible for maintaining the electronegativity of the entire tumor, primarily within it, during its growth over time. It is concluded that the anomalous bioelectricity and biomechanics in unperturbed cancer are due to intratumoral electrical heterogeneity and anisotropy according to Φ_1_(r, *θ, φ*), *ρ*_*v*_(*t*), *σ*_12_ and Turing spatiotemporal patterns, depending on the degree of *Y*_*nm*_ asymmetry, *µ*_*T*_ value 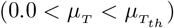 and *t*. Anomalous bioelectricity and biomechanics are closely related and both should be considered as another hallmark of cancer because these are essentially involved in self-regulation of symmetry, global electronegativity, intratumoral biological heterogeneity and anisotropy, growth, metastasis (by electrostatic and/or electromagnetic repulsion), abnormal metabolism of unperturbed cancer, as well as its protection from attack by cellular elements of the immune system and anticancer therapies (by formation of a heterogeneous electric shield), and in the appropriate and individualized selection of anticancer therapy, either alone or in combination.

**Author summary:** Unperturbed cancer and its growth kinetics are not fully understood, an aspect that may partially explain why its complete cure has not yet been achieved by any existing anticancer therapies. This may be because the close electrical-mechanical-chemical-biological-kinetics-hallmarks connection in unperturbed cancer is not well understood. Therefore, this study proposes a theoretical approach that reveals the explicit connection between anomalous bioelectricity (electrical properties of cancer and surrounding healthy tissue, electrical biopotentials, surface charge density and volumetric charge density) with mechanical (phase-field, cancer asymmetry degree and diffusive interface), chemical-biological (Turing approach) and kinetic (size, geometry and introduction of mass) parameters in unperturbed cancer. Simulations reveal: 1) this electrical-mechanical-chemical-biological-kinetics-hallmarks connection may be essentially governed by both electronegative and electropositive intratumoral regions, which are closely interconnected and dependent on spherical harmonic asymmetry degree and introduction of mass (less than its threshold value) during unperturbed cancer growth. 2) Greater spherical harmonic asymmetry and amount of mass introduced result in a greater degree of heterogeneity and faster growth of it. 3) Electronegative and electropositive intratumoral regions may be responsible for the self-preservation and self-regulation overtime of symmetry of electric potential and volumetric charge density spatial patterns, both global electronegativity and survival of unperturbed cancer.

## Introduction

The cancer is the second cause of death in worldwide and a serious health problem and challenge for the international scientific community. The genesis, growth, invasion and metastasis of unperturbed cancer and its protection against the immune system are not fully understood, despite unquestionable advances in molecular biology and genetics [1, 2], immunology in cancer [3], and quantum biology [4]. Several hallmarks of undisturbed cancer are involved in growth, metastasis, and protection against attack by external agents (e.g., immune system and anticancer therapies) [5]. Nevertheless, these hallmarks in cancer are not fully understood too, as intratumoral biological heterogeneity and anisotropy.

Intratumoral heterogeneity and anisotropy in unperturbed cancer have been addressed not only from biology [5, 6], but also from mechanical [7–12] and electrical [13] points of view. The mechanical and electrical properties of the unperturbed cancer and surrounding healthy tissue are closely related to physiological, biophysical and metabolic processes in the cancer [13–18]. Nevertheless, both physical properties require a better understanding to fully know their roles in genesis, growth, invasion and metastasis of unperturbed cancer, as well as its protection against attack by external agents.

Castañeda et al. [8] use the Montijo-Bergues-Bory-Gompertz (MBBG) equation and report theoretically and experimentally (in Ehrlich and fibrosarcoma Sa-37 unperturbed malignant tumors) that the fractal dimension of its contour/surface is less than 1 (disconnected contour/surface), which means the existence of pores/holes/tunnels in this contour/surface (porous contour/surface from biophysical point of view). Furthermore, they suggest that the endogenous angiogenesis of unperturbed cancer is an emerging process because the amount of new blood vessels formed depends on these pores/holes/tunnels, in agreement with González et al. [7]. It is well known the endogenous cancer angiogenesis is related to its growth, aggressiveness, invasion, metastasis, and protection against attack by external agents [19]. Nevertheless, the hallmarks of cancer, such as angiogenesis, are not only involved in these processes, but also the physical properties (e.g., electrical conductivity, permittivity and impedance) and physical magnitudes (e.g., electrical biopotentials, electric field, charge density, and both faradic and ionic current densities of the unperturbed cancer and the surrounding healthy tissue) [13]. Both physical properties and physical magnitudes constitute the bioelectricity of a normal or pathological tissue. This tissue bioelectricity is associated with its metabolism [15, 16, 18] and biophysical-chemical environment [20, 21].

Bioelectricity and metabolism differ markedly between normal and pathological tissues. The bioelectricity and metabolism of unperturbed cancer as a whole (both intratumoral regions and at contour/surface) are anomalous when compared to those of the surrounding normal tissue [15–18, 22–25].

The role of anomalous bioelectricity in unperturbed cancer has been reported in few studies [13, 24]. Its explicit connection with metabolism, tumor growth kinetics (TGK) and hallmarks of cancer has not been established, except in a previous theoretical study that proposes a approximate physical model to connect this anomalous bioelectricity with unperturbed cancer growth [13]. Bory-Prevez et al. [13] report for the first time an approximate analytical expression to calculate surface charge density (*σ*_12_) at Σ during the unperturbed spherical cancer growth. This expression of *σ*_12_ at Σ depends on geometric (tumor radius, *R*_*T*_ ) and bioelectric (endogenous electrical biopotential at the center, Φ_*o*_; endogenous electrical biopotential at Σ, Φ_*s*_; electrical conductivity, *η*_1_; and electrical permittivity, *ε*_1_) parameters of the unperturbed malignant solid tumor, as well as bioelectric parameters of the surrounding healthy tissue (electrical conductivity, *η*_2_; and electrical permittivity, *ε*_2_). For this, they [13] assume the existence of an electromotive force field, named **E**_*f*_, which represents the active bioelectricity in cancer due to the endogenous negative electrical biopotentials in it, marked in its center, as reported by Miklavčič et al. [26].

Bory-Prevez et al. [13] consider, in a first approximation, that endogenous negative electrical biopotentials in unperturbed cancer depend only on r (position of any point (x,y,z) inside to unperturbed spherical malignant tumor). Nevertheless, their theoretical approach does not include the following aspects: 1) angular coordinates of the spherical harmonics *θ* (zenith/polar angle) and *φ* (azimuthal angle). 2) Local changes, curvature and redistribution of negatively and positively charged carriers at Σ. 3) The term that allows describing the migration of negatively charged carriers from Σ into unperturbed cancer. 4) The explicit connection *σ*_12_ at Σ with the growth, progression, metastasis and aggressiveness of the solid tumor, as well as its protection against attack by external agents. 5) The term that may relate *σ*_12_ at Σ to angiogenesis [27], the fractal dimensions of the contour (*d*_*f*_ ) and mass (*D*_*f*_ ) of the unperturbed solid cancer, and nucleation and impingement mechanisms involve in unperturbed tumor growth kinetics (TGK) [7, 8, 28]. These aspects and others constitute limitations of the study reported by Bory-Prevez et al. [13].

Changes on the surfaces of biological membranes and tissues are not only influenced by *σ*_12_ [13], but also by their mechanical-chemical properties [29–37]. This may suggest that mechanical, electrical, and chemical properties are closely related to each other; and may influence the complete TGK of undisturbed cancer and all its inherent biophysical-chemical processes. Nevertheless, an experimental and/or theoretical study that integrates these three types of properties with each other in unperturbed cancer and their influences on TGK and biophysical-chemical processes inherent in it; metastasis; and protection against attack by external agents has not been reported in the literature, up to now. Therefore, the aim of this study is to simulate the mechanical-electrical coevolution of unperturbed cancer using an energetic approach, which involves the local spatiotemporal changes of anomalous bioelectricity in the unperturbed cancer that are related to the global anomalous bioelectricity of it. Both local and global changes influence the anomalous metabolism, complete TGK, intratumoral heterogeneity and anisotropy, phase transition, angiogenesis, and metastasis involve in unperturbed cancer, as well as in its protection against attack by external agents.

## Materials and methods

### Assumptions

1. A three-dimensional, conductive, anisotropic and heterogeneous region was formed by two linear, anisotropic and heterogeneous media (tumor and the surrounding healthy tissue) separated by an interface Σ. The untreated solid tumor (medium inside Σ, named medium 1) was considered as a heterogeneous conducting sphere of radius *R*_*T*_ (in m) of average electrical conductivity (*η*_1_, in S/m) and average electrical permittivity (*ε*_1_, in F/m). The surrounding healthy tissue (medium outside Σ, named medium 2) was supposed to be a heterogeneous infinite medium of average electrical conductivity (*η*_2_, in S/m) and average electrical permittivity (*ε*_2_, in F/m), where *η*_1_ *> η*_2_ and *ε*_1_ *> ε*_2_ (Fig 1).
2. The source of electricity was neglected because the tumor was unperturbed.
3. The Maxwell-Wagner-Sillars effect occured physiologically between the tumor and the surrounding healthy tissue (see Introduction section in [13]).
4. As a first approximation, **E**_*f*_ depended only on the distance to the tumor center.
5. Normal and cancer cells that were at Σ do not significantly contribute to **E**_*f*_ .
6. The reactive and diffusion effects of substances inside the tumor modified *σ*_12_ at Σ and electrical biopotentials both inside and on Σ of the solid tumor.
7. The electrical properties of this biological tissue generated important local changes in the bulk and surface of the tumor.
8. The growth of the tumor was modeled from the energetic approach using an order parameter and important feedback between growth and mechanical stresses through the redistribution of charge densities in the bulk of the tumor and on its surface.

### Further comments about assumptions

The assumptions 1-5 were reported in [13], whereas the assumptions 6-8 were based in previous studies [29–36]. In this study, the justifications for each of assumptions 1-5 were not included because they were also reported in c1, unlike assumptions 6-8. That is why assumptions 6-8 were only discussed and justified in this study.

**Fig 1.**
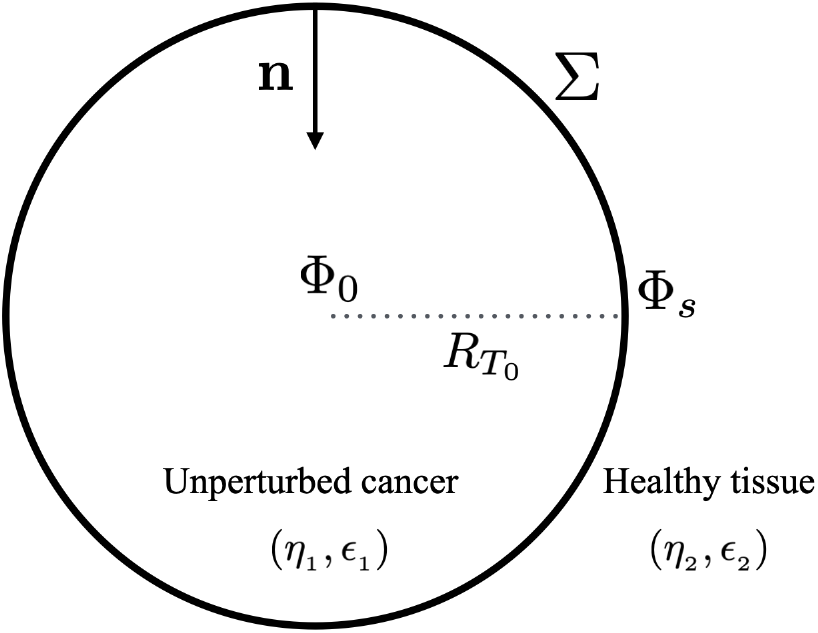
Spherical tumor at t = 0 days. Variables Φ_0_ and Φ_*s*_ denoted the electrical potentials in the center and the periphery of the tumor, respectively. 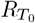 was the initial tumor radius. *η*_*i*_ and *ε*_*i*_ represented the electrical conductivities and electrical permittivities of the tumor (i = 1) and the surrounding healthy tissue (i = 2). **n** denoted the inward unit normal vector to the boundary Σ (interface that delimites both tissues).

The validation of assumptions 6-8 was demonstrated from mathematical and biological points of view in [29–31]. Ferre-Torres et al. [29] reported endogenous angiogenesis from an elastic mechanical point of view by means of the use of an energy functional to model the dynamics and morphology of sprouting. This energy functional considered effects of the extracellular matrix and the chemotactic factor in an order parameter that was locally modified on the surface by a spontaneous curvature effect. Likewise, Rueda-Contreras et al. [30] documented effects of pattern formation in growing domains with variable curvature for the same approximation of the order parameter. Furthermore, they demonstrated that the dynamics of some chemicals, modeled with a reaction-diffusion system, produced a three-dimensional pattern of chemical concentrations that modified the domain as it grew and this, in turn, affected the local mechanical stress in the domain [30].

Romero-Arias et al. [31] followed the same methodology of Rueda-Contreras et al. [30] to describe the viscoelastic properties on the surfaces of water droplets that contained electrical complexes on the surface. For this, they considered that the electrical effects of the dipole-dipole interaction were sufficient to determine the surface tension and shape of the drops. Therefore, we considered that the properties of surface charge density in tumors, nucleation, and angiogenesis could be addressed through the same formalism of the order parameter. Thus, we thought that local changes in the distribution of surface charges could be a reflection of the mechanical-electrical interactions on the surface. Consequently, the growth, progression, and metastasis of a solid tumor could be spatially and electrically correlated.

### Theory

The assumptions mentioned above and the close relationship between the physical and biological aspects in the unperturbed cancer allows to consider that Φ and **E**_*f*_ were related, in a first approximation, by means of the equation

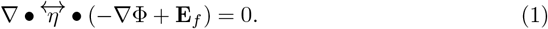

Due to the properties of the tensor 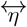 (real symmetric second-order tensors (3 x 3 symmetric matrix)), Bory-Prevez et al. [13] obtained

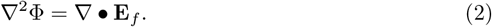

#### Boundary conditions

The region of interest is assumed to be a heterogeneous biological tissue formed by the unperturbed solid cancer (with average values *η*_1_ and *ε*_1_) surrounded by the surrounding healthy tissue (with average values *η*_2_ and *ε*_2_), as shown in Fig 1.

According to the continuity equation for the static case, the current density normal components of the tumor (**J**_1*n*_) and the surrounding healthy tissue (**J**_2*n*_) were continuous at Σ.

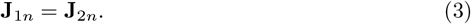

Therefore,

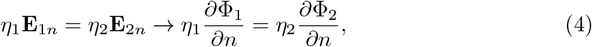

where **E**_1*n*_ is the normal component of the electrical field of unperturbed cancer. **E**_2*n*_ is the normal component of the electrical field of the surrounding healthy tissue. Φ_1_ is the electrical potential in unperturbed cancer and Φ_2_ the electrical potential in the surrounding healthy tissue. The normal derivatives of Φ_1_ and Φ_2_ are 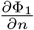 and 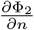, respectively.

Eq (3) was valid if **E**_*f*_ = **0** at Σ. The positive normal to unperturbed cancer surface was indicated as a unit vector **n** (represented schematically in Fig 1 by **n**) draw from the surrounding healthy tissue (medium 2) into unperturbed cancer (medium 1). According to this convention, medium 2 lay on the negative side (**n**_2_ = −**n**), and medium 1 on the positive side (**n**_1_ = **n**). Taking this into account the matching boundary condition for **D** (**D**_1*n*_ − **D**_2*n*_ = *σ*_12_), Eq (4) and **D** = *ε***E**yield the following expression for *ε*_12_,

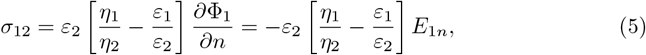

where *D*_1*n*_ and *D*_2*n*_ were the normal components of the flux density vector **D** in the tumor and the surrounding healthy tissue, respectively.

#### Calculation of the free electric charge surface density *σ*_12_

Strictly speaking, the problem to be solved for the calculation of the electric potential was Eq (2) subjected to the matching boundary conditions for Φ and 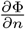

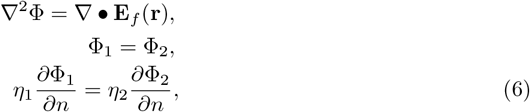

where **r** ∈ Σ.

The malignant tumor spherical model was reported in [11, 13, 38–41]. As Φ at Σ could be experimentally measured, conditions that could be replaced by a condition of Dirichlet and the work region was only inside the spherical malignant tumor, of radius *R*_*T*_, the solution of the problem of Poisson into the tumor in spherical coordinates (*r, θ, φ*; 0 ≤ *r < R*_*T*_, 0 ≤ *θ* ≤ *π*; 0 ≤ *φ* ≤ 2*π*) was given by

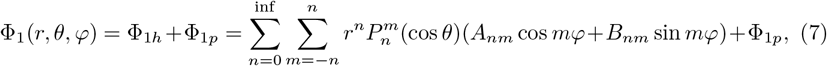

where 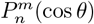 were the generalized Legendre polynomials order *n* and degree *m*. The Φ_1*p*_ was an any particular solution of Eq (2) in unperturbed cancer. The *A*_*nm*_ and *B*_*nm*_ were the coefficients of any homogeneous solution of Φ_1_(r, *θ, φ*).

#### Endogenous bioelectric potential

Eq (7) was rewritten as

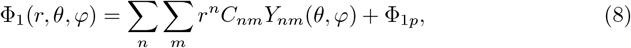

where *Y*_*nm*_ were the spherical harmonics with *C*_*nm*_ unknown. These harmonics were related to the generalized Legendre polynomials of by.

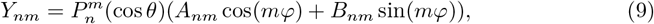

and the spherical harmonic complied with the following relationship

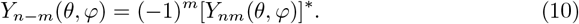

If we substitute *r* = 0 in Eq (8), we obtain Φ_1*p*_, given by

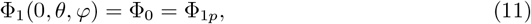

and Eq (8) becomes

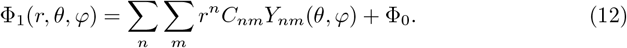

If the right hand of Eq (6) was zero (Laplace equation), its solution is Eq (9), where *C*_*nm*_

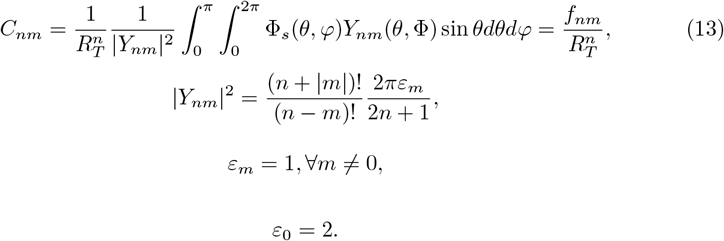

If we substitute *C*_*nm*_ (Eq (13)) in Eq (12), where Φ_*s*_(*θ, φ*) is the bioelectric potential on the surface of the unperturbed cancer, we obtain

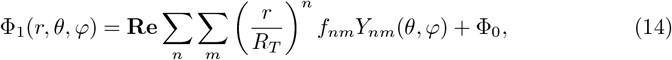

where **Re** signified the real part of the solution Eq (6).

Miklavčič et al. [26] measured endogenous bioelectric potentials from unperturbed cancer border to opposite along an axis (denominated z axis). In this case, Φ_1_(*z*, 0, 0) (*r* = *z, θ* = 0, *φ* = 0) is a parable, whose vertex coincided with unperturbed cancer center. Furthermore, they measured the bioelectric potential in Φ_1_(0, 0, 0) and 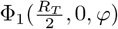 and their results showed that the bioelectric potential was more electronegative at unperturbed cancer center and less electronegative in its border. In other words, unperturbed cancer becomes less electronegative from the center towards Σ. The values of 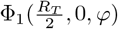 at *φ* = 45, 135, 225, 315^°^ were approximated and symmetrical.

#### Expression of the bioelectric potential for n = 2

As *f*_*nm*_ can be determined from experimental data of Φ_1_(r, *θ, φ*) [26], Eq (14) becomes

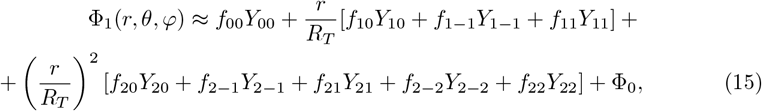

with

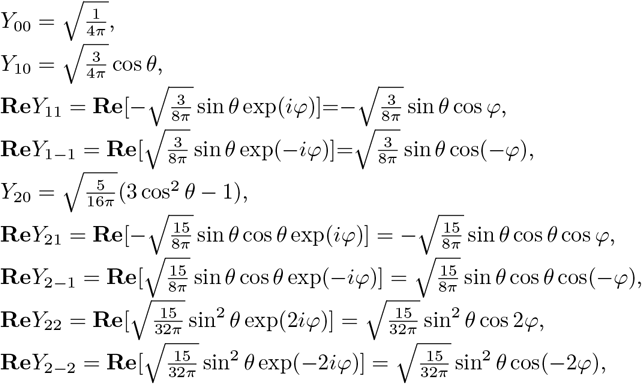

where *Y*_00_, *Y*_10_, **Re***Y*_11_, **Re***Y*_1−1_, *Y*_20_, **Re***Y*_21_, **Re***Y*_2−1_, **Re***Y*_22_, and **Re***Y*_2−2_ were the real part of the bioelectric potential. In this study, the real part of all spherical harmonics of any order *n* and degree *m* (**Re**[*Y*_*n*−*m*_]) are denoted *Y*_*nm*_.

If we take into account Eq (10), the real part of Φ_1_(r, *θ, φ*) and the spherical harmonics above, Eq (15) can be written as Eq (16)

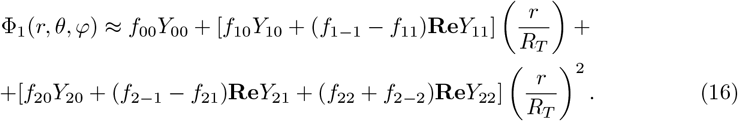

Since the unperturbed cancer was assumed to be spherical, the normal vector to the sphere is outward, and the normal derivative is

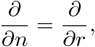

and from Eq (16), we obtained

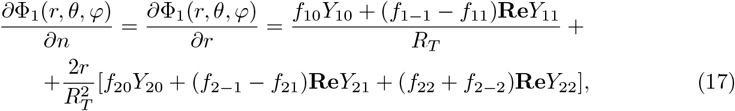

where 0 ≤ *r* ≤ *R*_*T*_, 0 ≤ *θ* ≤ *π*, 0 ≤ *φ* ≤ 2*π*.

The surface charge density was given by

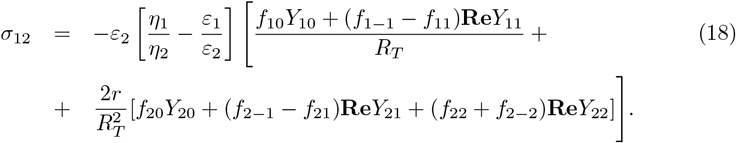

#### Radial dependence of Φ_1_(r, *θ, φ*)

If we neglect the quadratic term in Eq (16), Φ_1_(r, *θ, φ*) is written as

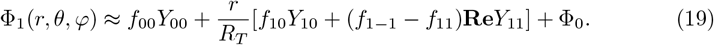

If Φ_1_(r, *θ, φ*) depended only on *r* (*θ* = 0 and *φ* = 0) in Eq (19), named Φ_1_(r), and compared it with Eq (2.10) reported by Bory-Prevez et al. [13], we obtain

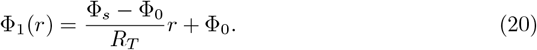

Thus,

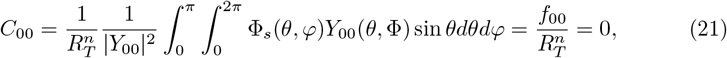

and

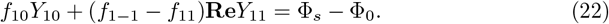

#### Energetic approach

The assumptions 6-8 may suggested that the chemical substances inside unperturbed cancer modify *σ*_12_ and the electrical biopotentials. On a simple form, reaction-diffusion processes for chemical substance with local curvature and mechanical-electrical interaction in a three-dimensional domain could be used to illustrate that the concentration of chemical substances modified the growth regions and the charge density in the tissue and, in turn, the redistribution of charge density modified the regions of growth and the electric patterns in unperturbed cancers.

An energetic approach of the phase field (PF) model, originally proposed to study the interface between two simple liquids, developed for non-equilibrium systems, had achieved great success in recent decades in describing a wide range of materials science and domain growth phenomena [30, 31, 34, 42–44]. PF model allows for an elegant and multifaceted numerical description of complex nonlinear problems with moving boundaries. Particularly, PF model is an adaptable method that can quantitatively describe a wide range of mechanical and dynamic properties of interfaces based on bulk properties. Furthermore, PF models require fewer parameters, making them suitable for modeling the morphology and growth of biological systems such as cell movement [45, 46], phyllotaxis [30], angiogenesis [29, 47, 48] and solid malignant tumor growth [47, 49].

The PF model consists of the free energy ℱ of the system in powers of a smooth scalar field (*ϕ* : Ω ⊂ **R**^**3**^ → **R**), which act as an order parameter

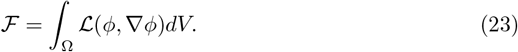

For an isotropic and homogeneous system, the order parameter represents two stable phases. Thus, the energy density ℒ could be written as 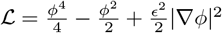, where the parameter *ϵ* represents a diffusive interface between the two phases [34]. One of the stable phases corresponded to the interior of the volume (*ϕ* = 1) delimited by unperturbed cancer surface located at *ϕ* = 0, another phase *ϕ* = − 1 represented the outer environment (surrounding healthy tissue).

We assume that electrical energy depended on the charge distribution and electrical potential, and in turn, the electrical potential depends on *σ*_12_, as is showed in Eq (18). Thus, we propose the term *B*_*ϕ*_ = *β*_0_(*ϕ*^2^ − 1)*σ*_12_ in the energy density, where *β*_0_ is a constant parameter that includes local modification of the surface in terms of a spontaneous change in curvature as a function of the surface charge density and endogenous potential, one sets 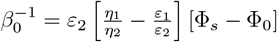. This assumption is focused on that *σ*_12_ at Σ can be produced an anisotropic bioelectrical effect in unperturbed cancer.

To mimic non-homogeneous growths on unperturbed cancer surface, we assumed that *σ*_12_ generates a mechanical energy cost the unperturbed cancer surface where the accumulation of charges at the surface constrained the mechanical structure of unperturbed cancer and conversely the surface structure of unperturbed cancer redirect *σ*_12_. Thus, mechanical-electrical term in the energy density was related with ∇*ϕ* ·∇*B*_*ϕ*_. Therefore, the total energy density was given by

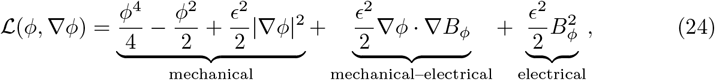

where the parameter *ϵ* represents a diffuse interface.

In this approach, the functional variations of the free energy with respect to the order parameter *ϕ* must then be subjected to diffusion, yielding a dynamic equation for the phase-field governed by 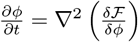, which could be expressed in terms of the order parameter and its spatial variations only [30, 32, 33, 43, 44, 50]. Nevertheless, the accumulation of cells inner unperturbed cancer must change this dynamics. Thereby, we proposed the evolution of *ϕ* as

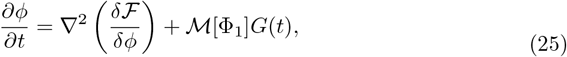

where 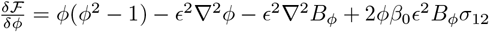. The functional ℳ [Φ_1_] represents the aggregation of masses in the inner phase *ϕ* = 1, which mimics the dependence of cancer cells on their endogenous biopotential. *G*[*t*] represents the tumor growth and its expression is given by the MBBG equation as [8]

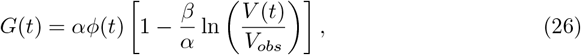

where V(t) representes the tumor volume at a time t after tumor cells are inoculated into the host. *V*_*obs*_ was considered as the tumor volume observed for the first time at latency time and it is palpable, observable but not measurable. *V*_*obs*_ had been estimated from experimental data. The parameter *α* is the intrinsic growth rate of the tumor. The parameter *β* is the growth deceleration factor due to endogenous antiangiogenic process [18, 51]. The solution of MBBG equation (Eq (26)) is given by

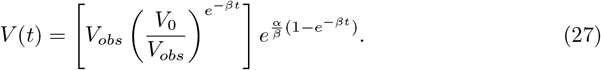

where the initial volume of unperturbed cancer satisfied the condition *V* (*t* = 0) = *V*_0_. For spheroidal tumors, 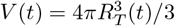 where *R*_*T*_ (*t*) is the radius of the tumor at a time *t*. As *R*_*T*_ (*t*) and *V* (*t*) depend on *t, R* in Eq (18) is replaced by *R*_*T*_ (*t*). As a result, *σ*_12_ is a function of *t*, namely *σ*_12_(*t*).

#### Chemical model

The evolution of a highly complex organism from a single cell or several micro-clusters of cells is a deeply intriguing subject, as suggested in unperturbed cancer [7, 52]. Hence, studying cancer development through the lens of morphogenesis can be particularly compelling. Mathematical and, more recently, computational studies of morphogenesis have shown that the essence of pattern formation was generally a complex developmental process [30, 42–44, 53–58]. Whereas early mathematical approaches on simple diffusion gradients and biochemical reactions, the advent of unprecedented computing power and its widespread accessibility have enabled increasingly sophisticated and realistic simulation studies. In particular, reaction-diffusion models were uniquely suited to spatially and functionally represent the complexity of a system that emerged collectively from a multiplicity of relatively simple signals, with a special emphasis on cell proliferation, migration and differentiation.

Morphogenesis explained by a reaction-diffusion mechanism with Turing system approach, is widely accepted. Under the right conditions, a reaction system involving two substances could produce periodic patterns through a diffusion-driven instability. Specifically, a rapidly diffusing global inhibitor interacts with a slowly diffusing local activator. Their functional coupling exhibits nonlinear reaction dynamics capable of generating repetitive patterns, such as spots or stripes [53, 57, 59, 60]. The patterns produced by gradients of inhibitors and activators, could be considered chemical pre-patterns that acts as templates for future differentiation and heterogeneous evolution on cancer cells. Thus, the apparent initial homogeneity of a cell group is transformed into spatially distinct profiles of regions with high and low concentrations that guide cell fate decisions and the emergence of visible forms.

Here, we propose a reaction-diffusion system as a chemical model (Turing patterns), which is solved to feed the phase-field with an initial condition. The mechanical formulation it was assumed that a set of cancer cells *u* compete with other set of cancer cells *v* to obtain space to proliferate. The first set of cancer cells are taken into account in the reaction-diffusion system, but they did not play a secondary role in the dynamics. The first set of cancer cells could be related to dormant/quiescent cells.

For the Turing dynamics, we adopted the Barrio–Varea–Aragon–Maini (BVAM) model [59, 61], which had proved to be applicable to model different biological processes. The BVAM model was written as

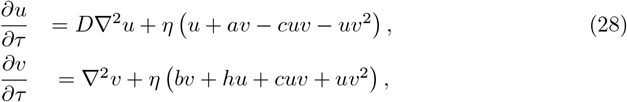

where, *D* was the ratio of the diffusion coefficients. The dimensionless constants *a, b, c* and *h* were kinetic parameters, and *η* gave the size of the system. The non-linear terms generated specify the pattern, the quadratic terms favors spots (*c* ≠ 0) and cubic terms favors stripes (*c* = 0). Here, we assumed that Turing process was faster than unperturbed cancer growth and deformation (*τ* ≫ *t*).

The BVAM model was applied in various biological systems [30, 61–66] and has the advantage of simulating several special functions on the sphere, such as the first spherical harmonics, or special functions that generate patterns in undefined geometric structures. Therefore, the BVAM model coupled with the phase-field model was a good combination for understanding tumor development in regular and irregular geometric structures.

In summary, this chemical model considered proliferating cancer cells as inhibitory agents that diffuse very rapidly in cancer tissue and, consequently, increases electrical activity within unperturbed cancer. Furthermore, this set of cancer cells generated spatial patterns in the volumetric charge distribution (*ρ*_*v*_(*t*)) that resembles the spatial functions given by *Y*_*nm*_ for a set of kinetic parameters. Thus, the chemical model coupled to the phase field allowed us to forget about the coordinate system, proposing a coordinate-free system where the boundary condition, given by *ϕ* = 0 and *σ*_12_, determined the electric biopotential inner unperturbed cancer. To show the relevance of the chemical and PF models in unperturbed cancer growth, we assumed that Φ_1_(*r, θ, φ*) followed the distribution of the inhibitor *v*(*τ* ) and we propose three ansatz

- The change in charge density throughout the interior of the unperturbed cancer volume over time is *dρ*_*v*_(*t*)*/dt* = *qv*(*t*),
- The surface charge density was 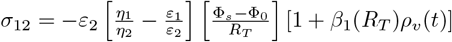,
- The mass aggregation on inner phase of *ϕ* is ℳ [Φ_1_] = *µ*_*T*_ *v*_+_(*t*),

where *κ* and *µ*_*T*_ are constants, *β*_1_(*R*_*T*_ ) is a parameter that depends on *R*_*T*_ and *v*_+_(*t*) are the positive values of *v*(*t*), *i*.*e*., there is only mass aggregation where the inhibitor is positive. From Eq (18), we assume *β*_1_(*R*_*T*_ ) = 2*/R*_*T*_ .

#### Stability of Turing patterns

If we consider that Eqs (28) is inserted into a unitary domain of size *L* = 1 and scale this dimension as *η* = *L*^2^*/aδ* for some scale factor *δ*. Then, the Turing instability of the stationary states (*u*_0_, *v*_0_) resulted from the calculation of the characteristic polynomial

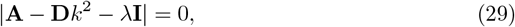

where *k* is the wave mode and

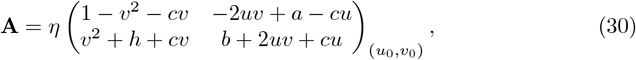

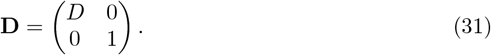

The unstable wave modes could be estimated by means of the critical value of the bifurcation parameter by noticing that at the onset of the instability *λ*(*k*_*c*_) = 0. Thus, the most unstable wave number when 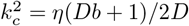 [59].

In the three-dimensional case, the nonlinear bifurcation analysis served to describe changes in the dynamics of the system around the linear stability when the parameters of the system varied. In a simple cubic lattice, the structures that arised from this analysis may get planar, cylindrical or spherical droplet arrangements for *k*_*c*_ = 0.85 only. This condition predicted that planar structures could be stable for *c <* 0.361, the spherical shapes were stable for 0.361 *< c <* 0.589, and the square packed cylinders were stable for all *c <* 0.650 [59, 61–63]. We aim to understanding pattern formation on a simulated sphere of radius, named *R*, for the purposes of our model. We quantified the effect of domain size on patterning. The linear stability analysis for a spherical domain yielded the range of wave numbers for which a pattern emerges. The rage of wave numbers satisfied the following condition [63]

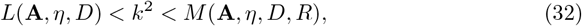

where the low interval was

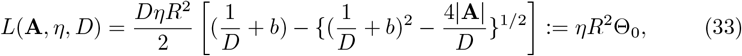

and the maximum interval was

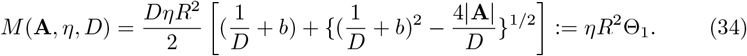

The associated Legendre equation showed that the eigenvalues *n* of *Y*_*nm*_ were related to the wave modes as *k*^2^ = *n*(*n* + 1) [63]. Therefore, for every *Y*_*nm*_ and radius *R*, we found a Turing pattern if we changed the scale factor *δ* as *η*_0_*R*^2^Θ_0_ *< δ* · *n*(*n* + 1) *< η*_0_*R*^2^Θ_1_, where 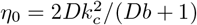. The values of *δ* were chosen for different symmetries considering that the wave modes *k*^2^ = *n*(*n* + 1) were the nearest possible to the ideal value for a given symmetry. Numerical simulations of *Y*_*nm*_ with the BVAM model for each *δ* on a sphere of radius *R* = 10 (unit in pixels) is showed in Fig 2. For this, we solved the equations of the chemical model (Eq (28)) for different values of *δ*. The symmetries on a sphere were highly correlated with the scale of the domain and also had an excellent matching with the reports of Turing patterns [62, 63].

**Fig 2.**
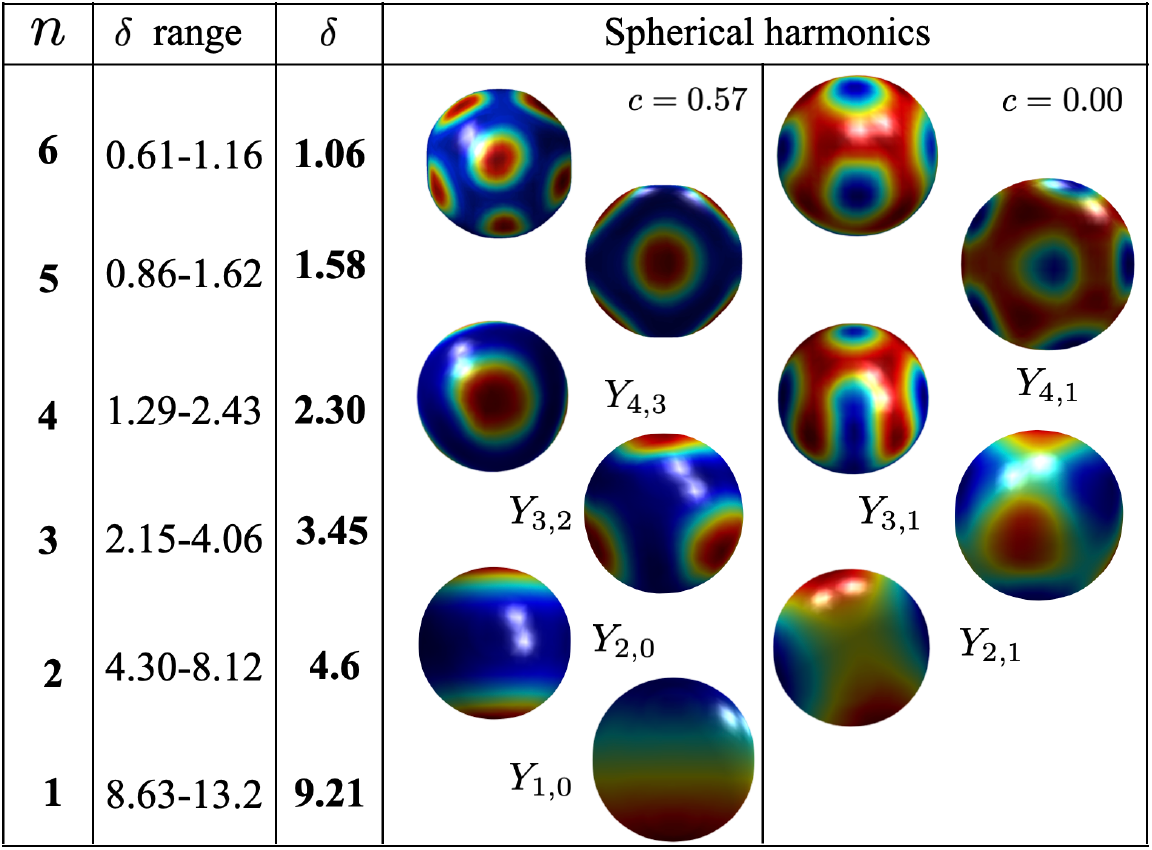
Spherical harmonics using Barrio–Varea–Aragon–Maini (BVAM) model for different spacial scales. Negative and positive of *ρ*_*v*_(*t*) represented inhibitor (in blue color) and activator (in red color) mechanisms of unperturbed cancer growth, respectively.

#### Numerical simulations

The physical magnitudes Φ_1_(r, *θ, φ*), *ρ*_*v*_(*t*) and *σ*_12_ were simulated in this study. The Φ_1_(r, *θ, φ*) was simulated from experimental and theoretical results reported in the literature [14, 25, 26, 38, 67, 68] and theoretically mimicked these results. The *σ*_12_ was considered in our simulations from theoretical results documented in [13] and generalized, taking into account spherical harmonics. The *ρ*_*v*_(*t*) was used because it was natural to examine the endogenous bioelectric potential of entire unperturbed cancer in terms of the Poisson equation and associate changes in tissue biomass with variations in volumetric charge density and vice versa.

We simulated *σ*_12_ versus *R*_*T*_ and *dσ*_12_*/dt* versus *σ*_12_ for three values of *CDR*/*CDR*_*PF*_ (1, 3 and 5); two values of Φ_0_ − Φ_*s*_ (- 145 and - 25 mV); and values of the model parameter showed in Table 1 to compare our results with those reported by Bory-Prevez et al. [13]. The *CDR* corresponds to the theoretical model (Eq (5)) reported in [13], whereas *CDR*_*PF*_ to the simulated model with the phase-field approach (PF) reported in this study (Eq (18)). The first terms of Eq (18) were used to calibrate phase–field model. Then, we studied the impact of *Y*_*nm*_ in charge distribution and unperturbed cancer growing. The parameter *ϵ*_2_ in Eqs (5) and (18) was calculated by the expression *ϵ*_2_ = *ϵ*_*r*2_*ϵ*_0_, where *ϵ*_0_ (8.85×10^12^ F/m) was the vacuum permittivity and *ϵ*_*r*2_ (4 × 10^7^) the relative permittivity of the muscle.

**Table 1.** Model parameter values.

| Name | Parameter | Values (PF) |
| --- | --- | --- |
| Grid size in 3D | $L^3$ | $33.6^3$ mm<sup>3</sup> ($60^3$ pixels<sup>3</sup>) |
| Diffuse interface | $\epsilon$ | 0.56 mm (1 pixel) |
| Initial unperturbed cancer radio | $R_{T_0}(R)$ | 5.6 mm (10 pixels) |
| Growth rate | $\alpha$ | $0.6$ days<sup>-1</sup> |
| Growth deceleration | $\beta$ | $0.2$ days<sup>-1</sup> |
| Inner electric potential | $\Phi_0$ | -160 mV |
| Surface electric potential | $\Phi_s$ | $[-15, -135]$ mV |
| Conductive–Dielectric ratio (CDR) | $[\eta_1/\eta_2 - \epsilon_1/\epsilon_2]$ | $[1, 5]$ |

In simulations, Φ_1_(r, *θ, φ*) was calculated using the Poisson equation and *σ*_12_ with the corresponding boundary conditions. The zero flux condition for PF was imposed in the cubic domain of integration, whereas the same condition was imposed in the boundary of unperturbed cancer (*ϕ* = 0) for Turing systems. The Φ_1_(r, *θ, φ*) had two Dirichlet conditions, one in the boundary of unperturbed cancer (-15 mV) and another in its center (-160 mV) when the Poisson equation was solved using *ρ*_*v*_(*t*) (charge density throughout the interior of the unperturbed cancer volume). The coefficients from Eq (13) were calculated for the initial condition and conserved throughout the simulation using *q* = 1.0. The increase of unperturbed cancer mass increased Φ_1_(r, *θ, φ*) following *Y*_*nm*_ in each time step.

In order to know how Φ_1_(r, *θ, φ*) and *ρ*_*v*_(*t*) space-time patterns in unperturbed cancer were influenced by *Y*_*nm*_ type and *µ*_*T*_, we used four values of *µ*_*T*_ (*µ*_*T*_ = 0.5, 1.0, 15.0 and 25.0) and four types of *Y*_*nm*_ (*Y*_20_, *Y*_21_, *Y*_32_ and *Y*_41_) at different time *t* (*t* = 0, 5, 10 and 15 days). Furthermore, animated movies for Φ_1_(r, *θ, φ*) and *ρ*_*v*_(*t*) were performed, and are shown in the Supporting Information.

Even (*Y*_20_) and odd (*Y*_21_, *Y*_32_ and *Y*_41_) spherical harmonics were used to understand how parity and higher-orders of *Y*_*nm*_ affect spatiotemporal distributions of these two physical quantities and contribute to biolectric heterogeneity. The *µ*_*T*_ values were used to mimic undifferentiated, moderately differentiated and differentiated/well-differentiated unperturbed cancer in clinics [69]. It is well documented that cells in undifferentiated unperturbed cancer do not resemble the normal cells of the tissue from which they originate; therefore, it grows more rapidly and are more aggressive. Cells in moderately differentiated unperturbed cancer show characteristics intermediate between normal and abnormal and by this reason it grows neither quickly nor slowly, and its aggressiveness is moderate. Cells in differentiated or well-differentiated unperturbed cancer resemble normal cells and tissues and consequently it grows more slowly and are less malignant [69].

Simulations mentioned in the previous paragraph allowed us to mimic the experimental results reported by Miklavčič et al. [26] (*µ*_*T*_ = 1.0 (reference value) and *t* = 0 days) and to know if those experimental results were valid for all *Y*_*nm*_, *µ*_*T*_ and *t*. Furthermore, both *Y*_*nm*_ and *µ*_*T*_ were used to understand how they influenced the unperturbed TGK (temporal behavior of *R*_*T*_ ) and the global bioelectricity (estimated with the norm of *ρ*_*v*_(*t*), named ||*ρ*_*v*_(*t*)||). The Φ_1_(r, *θ, φ*), *ρ*_*v*_(*t*) and *σ*_12_ spatiotemporal patterns were connected to *R*_*T*_ by means of ||*ρ*_*v*_(*t*)||.

In this study, the spherical harmonics *Y*_41_ (fourth-order odd harmonic) we used to understand how 2D Φ_1_(r, *θ, φ*) and *ρ*_*v*_(*t*) space-time patterns changed when the asymmetry of *Y*_*nm*_ increased (*Y*_41_ was more asymmetric than *Y*_32_). Furthermore, we used BVAM model to generate Turing patterns on unperturbed cancer spherical surface at *t* = 0, 5, 10 and 15 days and their corresponding *ρ*_*v*_(*t*) space-time patterns. For this, we used *δ* = 1.55 and *c* = 0.00. The Turing patterns represented unperturbed TGK with inhibitory mechanism (in blue with *ρ*_*v*_(*t*) *<* 0) and activating mechanism (in red with *ρ*_*v*_(*t*) *>* 0).

As Turing’s morphogenesis model is a fundamentally chemical-biological (reaction-diffusion) process and does not involve electrical charges or electrostatic principles, the two purposes of simulations of BVAM model were to know how these chemical-biological (reaction-diffusion) process were connected with *ρ*_*v*_(*t*) spatiotemporal patterns and to determine how Turing patterns mimicked dark and bright stripes observed in unperturbed cancer. This second purpose was based on the premise that the Turing pattern generated stripes (similar to those of a zebra or spots of a cheetah) due to areas/regions of high and low concentration of chemical substances, called morphogens, which interact, diffuse, and act as signals that indicate cells how to differentiate. These regions of high and low concentrations corresponded to those with high (activator processes) and low (inhibitor processes) intensity of the chemical signal from these morphogens, respectively. Furthermore, dark and bright stripes were observed by magnetic resonance imaging (MRI) and current density imaging (CDI) in Sa-1 sarcoma growing on male A/J mice (Fig 2) and T50/80 mammary carcinoma growing on immunodeficient mice (Fig 3) [14]. Serŝa *e*t. al [14] documented that dark stripes were observed in intratumoral regions of unperturbed cancer with low signal on MRI that correspond to high current density on CDI, whereas bright stripes were observed in those with high signal on MRI that corresponded to low current density on CDI.

**Fig 3.**
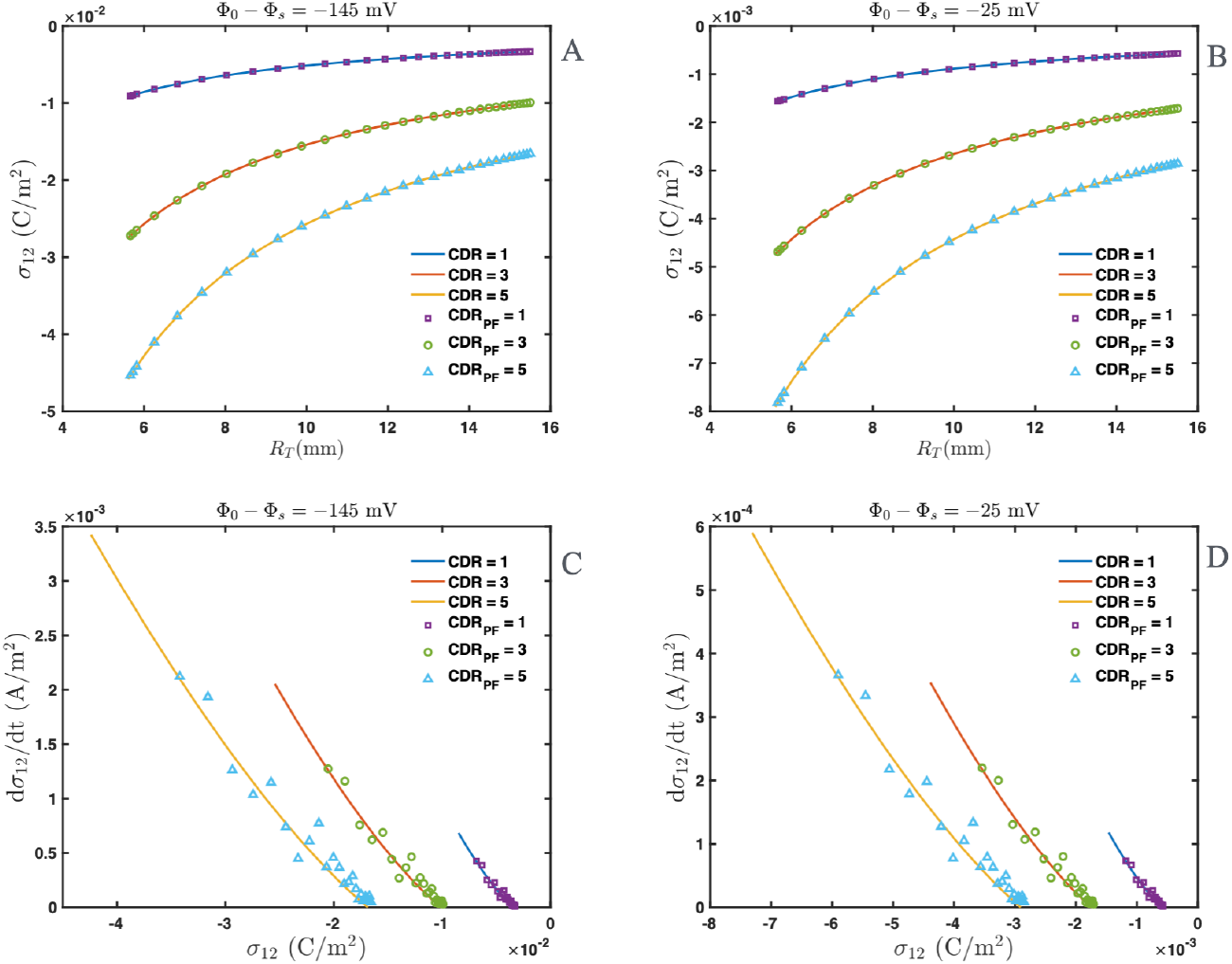
Surface chrage density and phase field model (PF) for different conductive-dielectric ratios (CDR). Free electric charge surface density (*σ*_12_) and and rate of change of *σ*_12_ over time (*dσ*_12_*/dt*) in unperturbed tumor for first order of Eq (18) for different values of *CDR*/*CDR*_*PF*_ and Φ_0_ − Φ_*s*_. (A) *σ*_12_ versus *R*_*T*_ (tumor radius) for Φ_0_ − Φ_*s*_ = - 145 mV. (B) *σ*_12_ versus *R*_*T*_ for Φ_0_ − Φ_*s*_ = - 25 mV. (C) *dσ*_12_*/dt* versus *σ*_12_ for Φ_0_ − Φ_*s*_ = - 145 mV. (D) *dσ*_12_*/dt* versus *σ*_12_ for Φ_0_ − Φ_*s*_ = - 25 mV. Values of *CDR*/*CDR*_*PF*_ = 1, 3 and 5 were used in these four subplots. The solid lines and different symbols represent values of Conductive-Dielectric ratio (*CDR*) theoretical model [13] and simulated models with the phase-field approach, PF (*CDR*_*PF*_ ), respectively.

The global value of *ρ*_*v*_(*t*) in unperturbed cancer volume was calculated by two different metrics (Euclidean norms in ℝ^*n*^), named ||*ρ*_*v*_(*t*)|| and ||*ρ*_*vs*_(*t*)|| norms, for spherical harmonics *Y*_20_, *Y*_21_ and *Y*_32_ and *µ*_*T*_ = 0.5, 1.0, 25.0 and 50.0 at each instant *t*. The first norm was calculated by means of 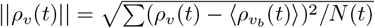, where *ρ*_*v*_(*t*) and ⟨*ρ*_*vb*_ (*t*) ⟩ were the volumetric charge densities on throughout unperturbed cancer (interior and surface) and intratumor spherical shell of width 2*ϵ*, respectively. The sum was over unperturbed cancer domain with size *N* (*t*). This spherical shell of width 2*ϵ* was spatially located between the surface of the unperturbed cancer and its intratumoral region that contained all negative and negative zones of *ρ*_*v*_(*t*). Consequently, ||*ρ*_*v*_(*t*)|| only sensed the global changes that occurred in all these negative and negative zones of *ρ*_*v*_(*t*). The second norm was computed using 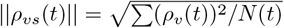 and included all values of *ρ*_*v*_(*t*) throughout the entire unperturbed cancer (interior and surface). Therefore, ||*ρ*_*vs*_(*t*)|| norm did not discriminate changes of *ρ*_*v*_(*t*) in negative and negative intratumor zones, spherical layer, and surface of unperturbed cancer. The || *ρ*_*v*_(*t*) || and ||*ρ*_*vs*_(*t*) || norms allowed us to understand whether the entire unperturbed TGK was influenced by local (intratumoral negative and negative zones) or global (entire volume) changes in unperturbed cancer bioelectricity. For this, we proposed the plots of *R*_*T*_ versus *t*, ||*ρ*_*v*_(*t*)|| versus *t*, ||*ρ*_*v*_(*t*)|| versus *R*_*T*_, ||*ρ*_*vs*_(*t*)|| versus *t*, and ||*ρ*_*vs*_(*t*)|| versus *R*_*T*_ for each spherical harmonics, four values of *µ*_*T*_ and *t*.

In order to rescale the size of the domain, we considered that Eq (28) was setting into a unitary domain of size *L* = 1 and scale the dimension as *η* = *L*^2^*/aδ* for some scale factor *δ*. This let us to find a reference values of *δ*. In our simulations, the value of *δ* = 5.18 was used for *Y*_20_ and *Y*_21_, *δ* = 2.59 for *Y*_32_ and *δ* = 1.49 for *Y*_41_. These values were rescaled to reproduce Turing patterns on a sphere and maintaining its range at a central level to ensure that the typical Turing instability preserved the pattern during the volume increment.

The parameters of the Turing model were chosen from references [30, 59, 62, 63] to obtain the instability: *a* = 1.1123, *b* = − 1.0122, *h* = − 1 and *D* = 0.516 for all simulations. Instability in unperturbed cancer has been explained from different noise sources, which cause fluctuations and stochasticity in unperturbed cancer and its TGK [70]. Furthermore, the parameter *c* was varied in the range 0 ≤ *c* ≤ 0.57 to obtain different types of patterns. The parameter *η* was first varied to obtain the distinct *Y*_*nm*_ in the sphere. Turing simulations were initialized with a random distribution of *u* and *v* as considering a perturbation around the steady state (0, 0) of the model. Different patterns depended on the values of *η* and *µ*_*T*_ . These parameters determined the influence of the surface structure of unperturbed cancer and the bioelectrical potential on the local curvature.

All partial differential equations were numerically solved with MATLAB and Simulink (version 2024b, the MathWorks, Inc., 2024). This software ran on a Macbook Pro 64-bits with Apple M4 Pro chip and macOS Sequoia 15.6.1, located at the Department of Mathematics and Mechanics at IIMAS, UNAM. We used standard second-order finite differences for the spatial dependence of phase field and Turing models. The forward Euler method was used for the time dependence [30] for all unperturbed cancer equations.

An initial domain of *ϕ* consists on a sphere of radius *R* = 10 pixels on a cubic grid of *Nx* = *Ny* = *Nz* = 60 pixels. Each pixel represented a real unit to ensure that the space step *dx* = 1 was equivalent to *R*_*T*0_ = 5.6 *mm*. Two time steps were used: *dt* = 1 × 10^− 5^ (for the evolution of Φ_1_(r, *θ, φ*) and *ρ*_*v*_(*t*)) and *dτ* = 5 × 10^−2^ (for the Turing model). This guaranteed that the Turing mechanism was faster than the phase-field mechanism and charge density distributions. For each time cycle, the PF model was integrated over 1000 time steps of *dt* whereas Turing systems was integrated over 60 time steps *dτ* . An unperturbed cancer of 15 days of growth with *µ*_*T*_ = 1.0 required 300 cycles, which means that *dt* was equivalent to 4.32 seconds.

## Results

The simulations of *R*_*T*_ versus *t* and *σ*_12_ versus *t* had similar behaviors when time elapsed. The final value of *R*_*T*_ corresponded to *t* = 40 days after inoculation of cancer cells into the host. Fig 3 shows the plots of *σ*_12_ versus *R*_*T*_ (Figs 3A,B) and *dσ*_12_*/dt* versus *σ*_12_ (Figs 3C,D) for three values of *CDR*/*CDR*_*PF*_ (1, 3 and 5) and two values of Φ_0_ − Φ_*s*_ (- 145 mV (Figs 3A,C) and - 25 mV (Figs 3B,D)). In addition, this Fig 3 showed the comparisons between plots of *σ*_12_ versus *R*_*T*_ when these three values of *CDR* (theoretical model) and *CDR*_*PF*_ (simulated models with the phase-field approach (PF)) are used for each fixed value of Φ_0_ − Φ_*s*_. Similar comparisons were made for the plot *dσ*_12_*/dt* versus *σ*_12_.

Figs 3A,B show that *σ*_12_ versus *R*_*T*_ growth asymptotically in time for all values of Φ_0_ − Φ_*s*_ and *CDR*/*CDR*_*PF*_ . In other words, the tumor electronegativity grew asymptotically as *R*_*T*_ increases for all values of Φ_0_ − Φ_*s*_ and *CDR*/*CDR*_*PF*_ . This variation in tumor electronegativity (or *σ*_12_) was marked for the largest difference of Φ_0_ − Φ_*s*_ and the highest value of *CDR*/*CDR*_*PF*_ . Furthermore, Figs 3C,D reveal that *dσ*_12_*/dt* decreased asymptotically when *σ*_12_ increase, which is marked for the largest difference of Φ_0_ − Φ_*s*_ and the highest value of *CDR*/*CDR*_*PF*_ . The largest value of *dσ*_12_*/dt* occurred for the most negative value of *σ*_12_. The value of *dσ*_12_*/dt* = 0 indicates that *σ*_12_ remains constant over time from the value of *σ*_12_ that intersected the x-axis.

In both plots *σ*_12_ versus *R*_*T*_ and *dσ*_12_*/dt* versus *σ*_12_, there was good correspondence between the curves obtained with the same value of *CDR* and *CDR*_*PF*_ for each value of Φ_0_ − Φ_*s*_ (Fig 3).

The results of the simulations of two-dimensional (2D) Φ_1_(r, *θ, φ*) (in the mid-z plane (cross-section) of the sphere of radius *R*_*T*_ (−*R*_*T*_ ≤ *z* ≤ *R*_*T*_ ) that changed over time) and three-dimensional (3D) *ρ*_*v*_(*t*) space-time patterns dependent on *µ*_*T*_ and *Y*_*nm*_ (single or combined) for each *t* were shown in Figs 4-11. These 3D space-time patterns of *ρ*_*v*_(*t*) were displayed in such a way that the observer was outside the unperturbed cancer. Furthermore, animated movies of Φ_1_(r, *θ, φ*) and *ρ*_*v*_(*t*) for Figs 4-11 were shown in their respective Supporting Information S1-S9 Videos. These movies were obtained by reconstructing the space-time patterns of these two physical quantities in each z plane (−*R*_*T*_ ≤ *z* ≤ *R*_*T*_ ). In all these simulations and animated movies at *t* = 0 days corresponded with the time instant when unperturbed cancer reached *R*_*T*_ = 5.6 mm in the experiment [26].

**Fig 4.**
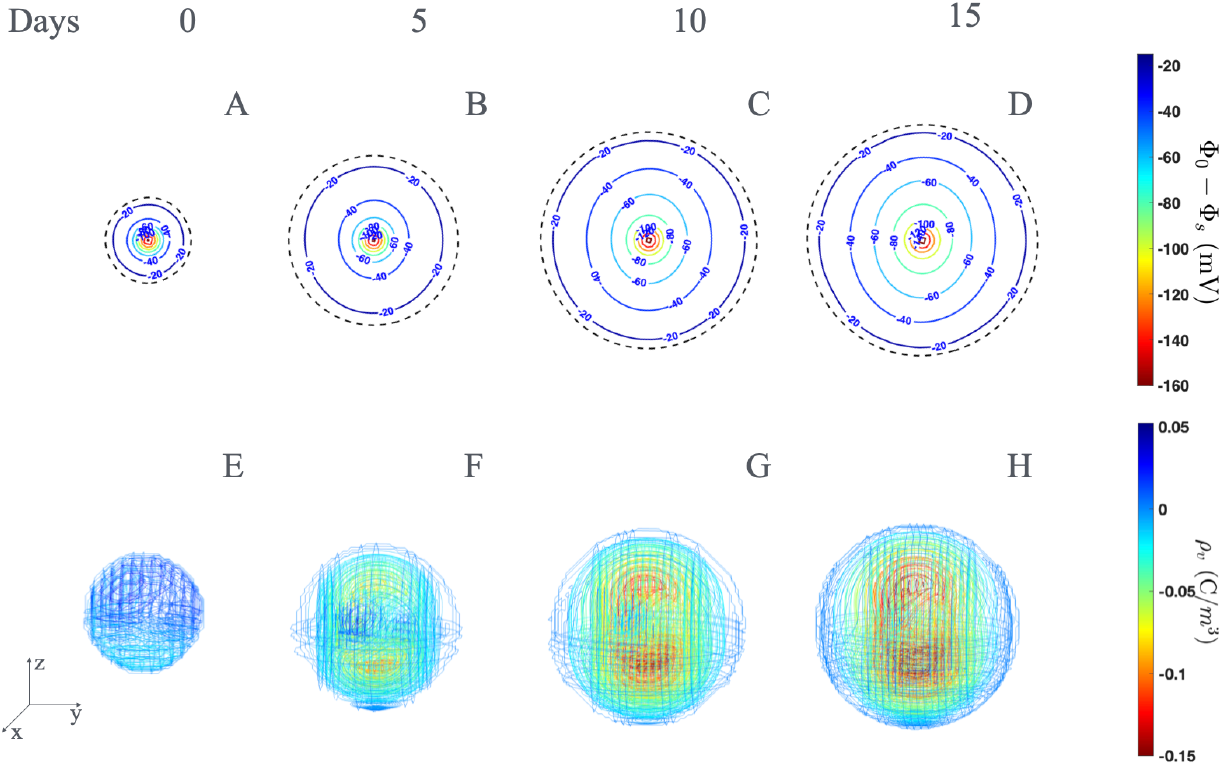
Biopotentials and volumetric charge density of unperturbed cancer for *Y*_2*m*_ spherical harmonic. Simulations of 2D electrical biopotentials Φ_1_(r, *θ, φ*) (A-D) and charge density *ρ*_*v*_(*t*) (E-H) spatiotemporal patterns obtained with Eqs (18) and (28). These patterns are showed at *t* = 0 days (A,E), *t* = 5 days (B,F), *t* = 10 days (C,G) and *t* = 15 days (D,H) for *µ*_*T*_ = 1.0.

**Fig 5.**
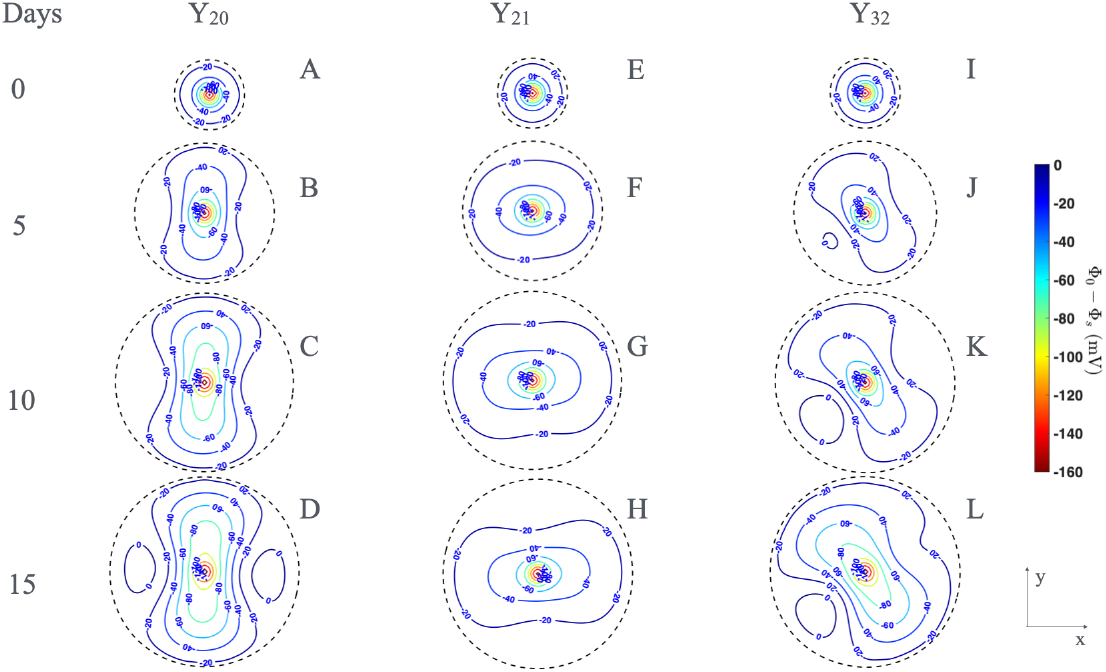
Biopotentials for different spherical harmonics. 2D space-time patterns (mid-z plane) of bioelectrical potentials (Φ_1_(r, *θ, φ*)) inside the unperturbed tumor for *µ*_*T*_ = 1.0 and three types of spherical harmonics at different times *t*. (A) *Y*_20_ at *t* = 0 days (B) *Y*_20_ at *t* = 5 days. (C) *Y*_20_ at *t* = 10 days. (D) *Y*_20_ at *t* = 15 days. (E) *Y*_21_ at *t* = 0 days. (F) *Y*_21_ at *t* = 5 days. (G) *Y*_21_ at *t* = 10 days. (H) *Y*_21_ at *t* = 15 days. (I) *Y*_32_ at *t* = 0 days. (J) *Y*_32_ at *t* = 5 days. (K) *Y*_32_ at *t* = 10 days. (L) *Y*_32_ at *t* = 15 days. The black dotted line represents the unperturbed tumor size during its growth at different times.

**Fig 6.**
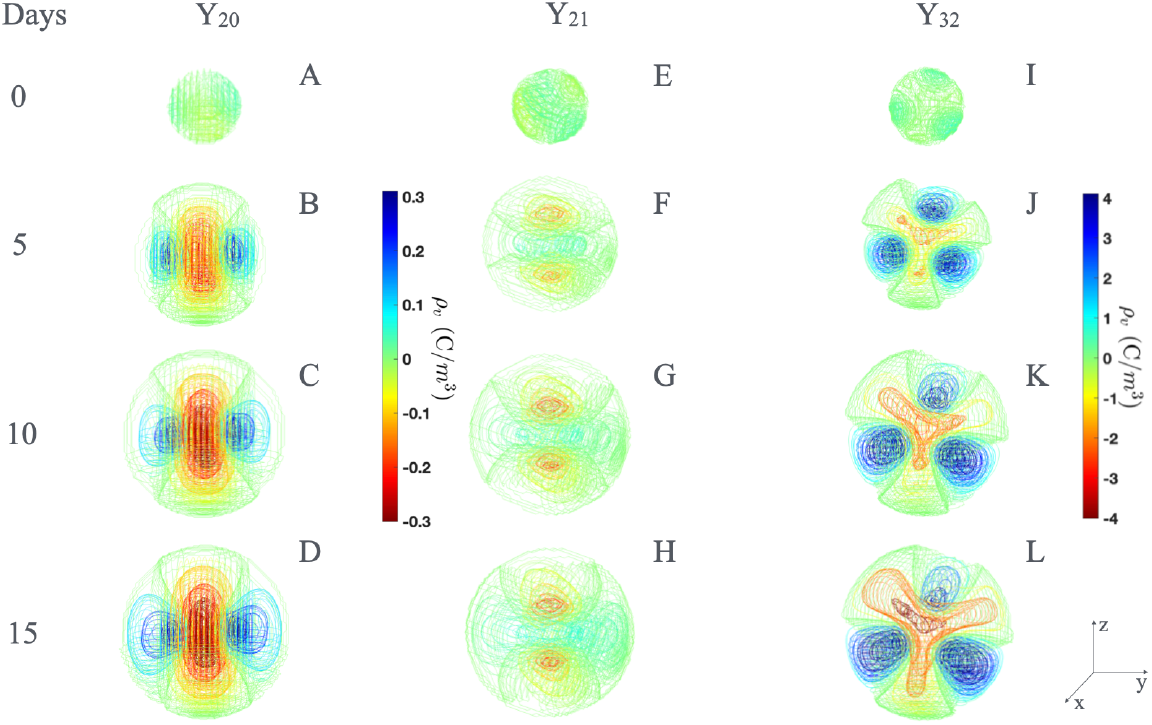
Volumetric charge density for different spherical harmonics. Space-time patterns of the volumetric charge density, *ρ*_*v*_(*t*), inside the unperturbed tumor for *µ*_*T*_ = 1.0 and three types of spherical harmonics at different times *t*. (A) *Y*_20_ at *t* = 0 days (B) *Y*_20_ at *t* = 5 days. (C) *Y*_20_ at *t* = 10 days. (D) *Y*_20_ at *t* = 15 days. (E) *Y*_21_ at *t* = 0 days. (F) *Y*_21_ at *t* = 5 days. (G) *Y*_21_ at *t* = 10 days. (H) *Y*_21_ at *t* = 15 days. (I) *Y*_32_ at *t* = 0 days. (J) *Y*_32_ at *t* = 5 days. (K) *Y*_32_ at *t* = 10 days. (L) *Y*_32_ at *t* = 15 days.

**Fig 7.**
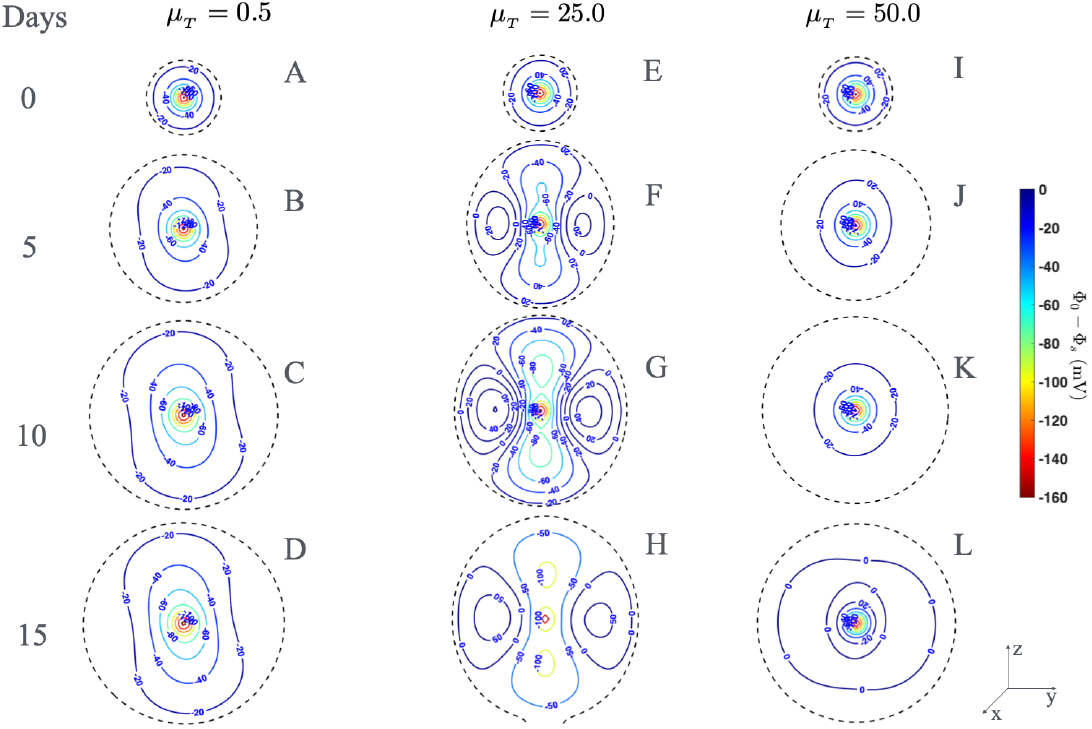
Biopotentials of *Y*_20_ for different *µ*_*T*_ values. 2D space-time patterns (mid-z plane) of bioelectrical potentials (Φ_1_(r, *θ, φ*)) inside the unperturbed tumor for three values of *µ*_*T*_, four values of time *t* and spherical harmonic *Y*_20_. (A) *µ*_*T*_ = 0.5 at *t* = 0 days (B) *µ*_*T*_ = 0.5 at *t* = 5 days. (C) *µ*_*T*_ = 0.5 at *t* = 10 days. (D) *µ*_*T*_ = 0.5 at *t* = 15 days. (E) *µ*_*T*_ = 25.0 at *t* = 0 days. (F) *µ*_*T*_ = 25.0 at *t* = 5 days. (G) *µ*_*T*_ = 25.0 at *t* = 10 days. (H) *µ*_*T*_ = 25.0 at *t* = 15 days. (I) *µ*_*T*_ = 50.0 at *t* = 0 days. (J) *µ*_*T*_ = 50.0 at *t* = 5 days. (K) *µ*_*T*_ = 50.0 at *t* = 10 days. (L) *µ*_*T*_ = 50.0 at *t* = 15 days. The black dotted line represents the unperturbed tumor size during its growth at different times.

**Fig 8.**
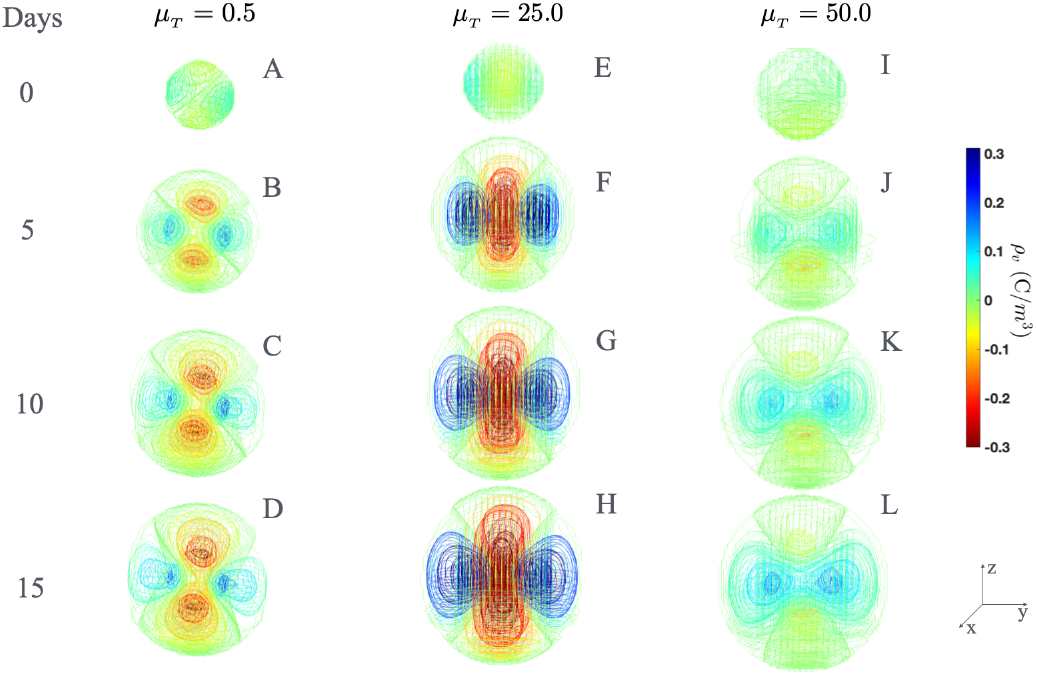
Volumetric charge density of *Y*_20_ for different *µ*_*T*_ values. Space-time patterns of the volumetric charge density, *ρ*_*v*_(*t*), inside the unperturbed tumor for three values of *µ*_*T*_, four values of time *t* and spherical harmonic *Y*_20_. (A) *µ*_*T*_ = 0.5 at *t* = 0 days (B) *µ*_*T*_ = 0.5 at *t* = 5 days. (C) *µ*_*T*_ = 0.5 at *t* = 10 days. (D) *µ*_*T*_ = 0.5 at *t* = 15 days. (E) *µ*_*T*_ = 25.0 at *t* = 0 days. (F) *µ*_*T*_ = 25.0 at *t* = 5 days. (G) *µ*_*T*_ = 25.0 at *t* = 10 days. (H) *µ*_*T*_ = 25.0 at *t* = 15 days. (I) *µ*_*T*_ = 50.0 at *t* = 0 days. (J) *µ*_*T*_ = 50.0 at *t* = 5 days. (K) *µ*_*T*_ = 50.0 at *t* = 10 days. (L) *µ*_*T*_ = 50.0 at *t* = 15 days.

**Fig 9.**
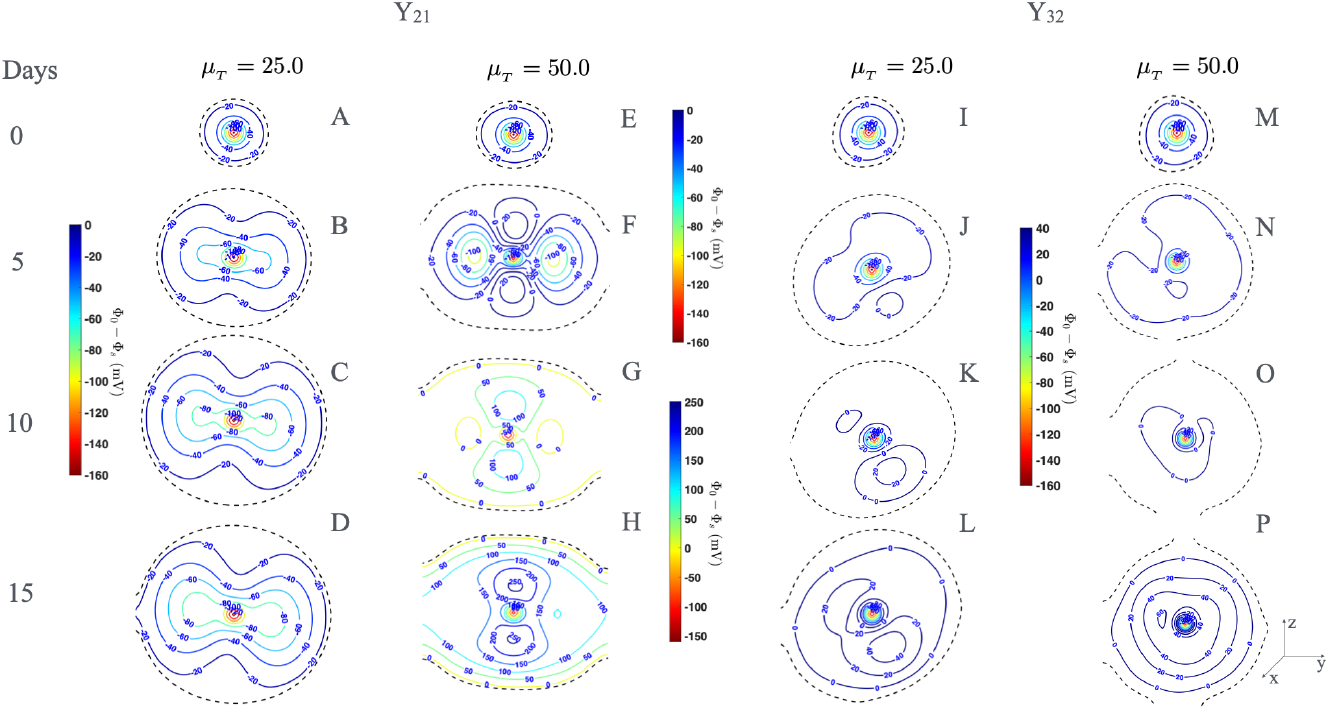
Biopotentials for different spherical harmonics and *µ*_*T*_ values. 2D space-time patterns (mid-z plane) of bioelectrical potentials (Φ_1_(r, *θ, φ*)) inside the unperturbed tumor for two values of *µ*_*T*_, four values of time *t* and spherical harmonics *Y*_21_ and *Y*_32_. (A) *µ*_*T*_ = 25.0 at *t* = 0 days. (B) *µ*_*T*_ = 25.0 at *t* = 5 days. (C) *µ*_*T*_ = 25.0 at *t* = 15 days. (D) *µ*_*T*_ = 25.0 at *t* = 15 days. (E) *µ*_*T*_ = 50.0 at *t* = 0 days. (F) *µ*_*T*_ = 50.0 at *t* = 5 days. (G) *µ*_*T*_ = 50.0 at *t* = 15 days. (H) *µ*_*T*_ = 50.0 at *t* = 15 days. (I) *µ*_*T*_ = 25.0 at *t* = 0 days. (J) *µ*_*T*_ = 25.0 at *t* = 5 days. (K) *µ*_*T*_ = 25.0 at *t* = 15 days. (L) *µ*_*T*_ = 25.0 at *t* = 15 days. (M) *µ*_*T*_ = 50.0 at *t* = 0 days. (N) *µ*_*T*_ = 50.0 at *t* = 5 days. (O) *µ*_*T*_ = 50.0 at *t* = 15 days. (P) *µ*_*T*_ = 50.0 at *t* = 15 days. The black dotted line represents the unperturbed tumor size during its growth at different times.

**Fig 10.**
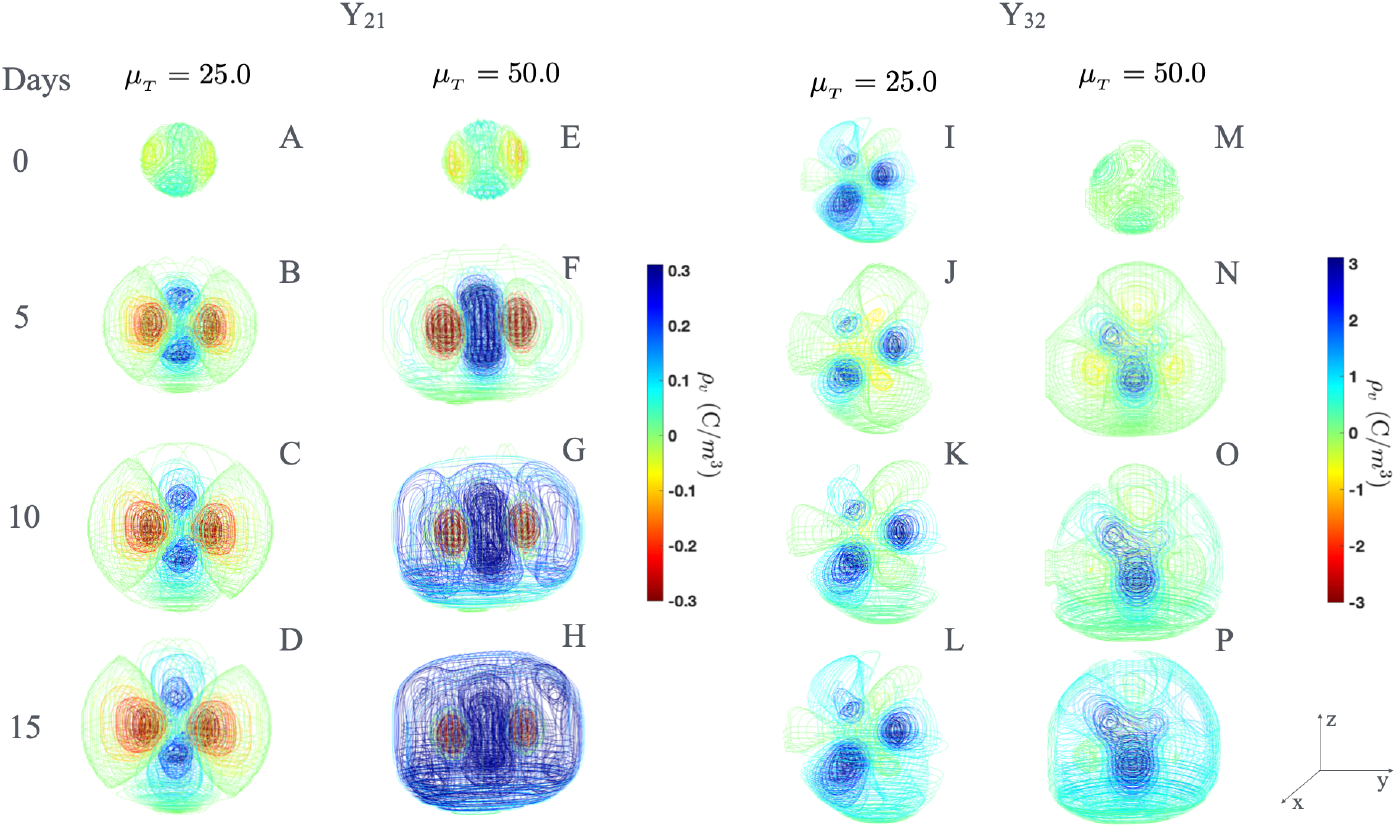
Volumetric charge density for different spherical harmonics and *µ*_*T*_ values. Space-time patterns of the volumetric charge density, *ρ*_*v*_(*t*), inside the unperturbed tumor for two values of *µ*_*T*_, four values of time *t* and spherical harmonics *Y*_21_ and *Y*_32_. (A) *µ*_*T*_ = 25.0 at *t* = 0 days. (B) *µ*_*T*_ = 25.0 at *t* = 5 days. (C) *µ*_*T*_ = 25.0 at *t* = 15 days. (D) *µ*_*T*_ = 25.0 at *t* = 15 days. (E) *µ*_*T*_ = 50.0 at *t* = 0 days. (F) *µ*_*T*_ = 50.0 at *t* = 5 days. (G) *µ*_*T*_ = 50.0 at *t* = 15 days. (H) *µ*_*T*_ = 50.0 at *t* = 15 days. (I) *µ*_*T*_ = 25.0 at *t* = 0 days. (J) *µ*_*T*_ = 25.0 at *t* = 5 days. (K) *µ*_*T*_ = 25.0 at *t* = 15 days. (L) *µ*_*T*_ = 25.0 at *t* = 15 days. (M) *µ*_*T*_ = 50.0 at *t* = 0 days. (N) *µ*_*T*_ = 50.0 at *t* = 5 days. (O) *µ*_*T*_ = 50.0 at *t* = 15 days. (P) *µ*_*T*_ = 50.0 at *t* = 15 days.

**Fig 11.**
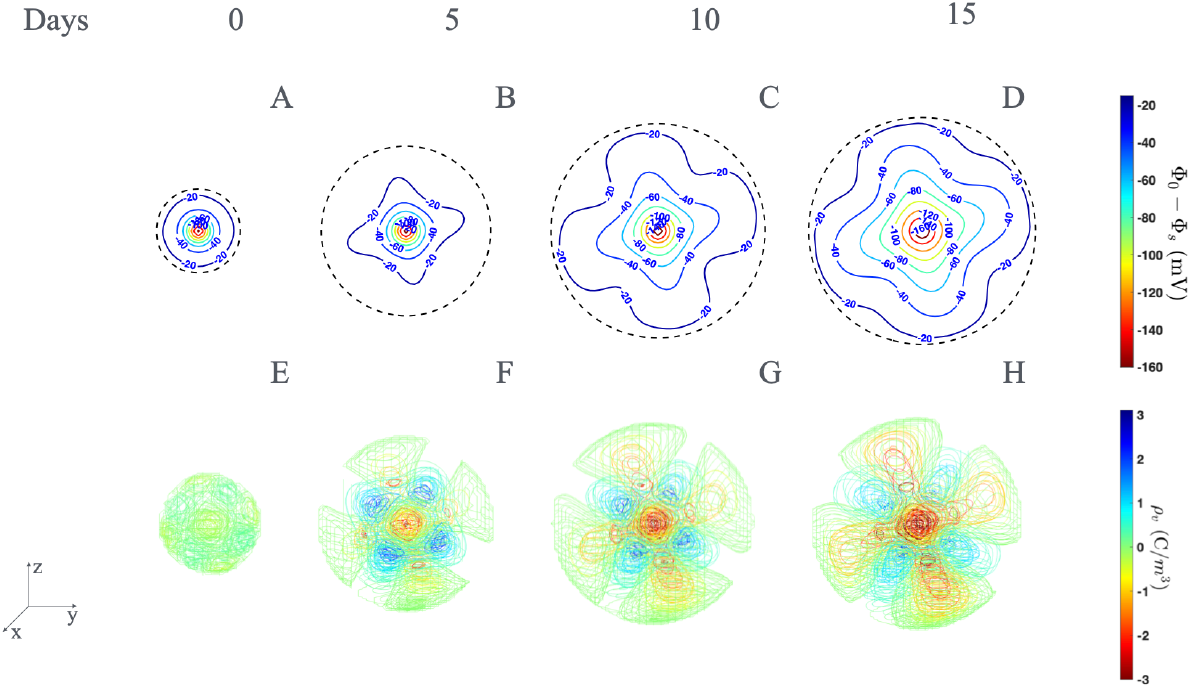
Biopotentials and volumetric density for *Y*_41_ spherical harmonic. Simulations of charge density *ρ*_*v*_(*t*) (Figs 4A-D) and 2D electrical biopotentials Φ_1_(r, *θ, φ*) (E-H) spatiotemporal patterns obtained for *Y*_41_. These patterns are showed at *t* = 0 days (A,E), *t* = 5 days (B,F), *t* = 10 days (C,G) and *t* = 15 days (D,H) for *µ*_*T*_ = 1.0 and *c* = 0.00.

Fig 4 showed 2D Φ_1_(r, *θ, φ*) (Figs 4A-D) and *ρ*_*v*_(*t*) (Figs 4E-H) space-time patterns for unperturbed cancer when Eqs (18) and (28) (two coupled chemical models) were solved. These 2D and 3D spatiotemporal patterns were depicted at *t* = 0 days (Figs 4A,E), 5 days (Fig 4B,F), 10 days (Fig 4C,G) and 15 days (Fig 4D,H). These figures were obtained for two values of *c* and *Y*_20_, *Y*_21_ and *Y*_32_. For *Y*_20_, *c* = 0.57 and the value of the inhibitory function *v*(*t*) for *σ*_12_ were taken. For *Y*_21_ and *Y*_22_, another chemical model with *c* = 0.000 was used and *σ*_12_ was overlapped on the values of the inhibitory functions *v*(*t*) for *Y*_21_ and *u*(*t*) for *Y*_22_. The values of the coefficients *f*_*nm*_ were estimated for an ellipsoid aligned along the principal axes starting from a slightly deformed sphere of radius *R*_*T*_ . We took *f*_00_ = 0, *f*_10_*Y*_10_ + (*f*_1−1_ − *f*_11_)**Re***Y*_11_ = Φ_*s*_ − Φ_0_, *f*_20_ = 1, (*f*_2−1_ - *f*_21_) = 0.5, and (*f*_22_ + *f*_2−2_) = 0.5. Animation movies of Φ_1_(r, *θ, φ*) and *ρ*_*v*_(*t*) based on the results in Fig 4 were shown in S1 Video in the Supporting Information.

The results depicted in Fig 4 and S1 Video Supporting Information revealed that *ρ*_*v*_(*t*) patterns changed in space-time by means of the self-organization of polarity throughout the unperturbed cancer volume: more negative polarity (central region), less negative polarity (inner non-central region) and weak positive polarity (spherical shell with a width 2*ϵ*, in which *ρ*_*v*_(*t*) → 0), as showed in Figs 4E-H. Despite this self-rearrangement of *ρ*_*v*_(*t*), the unperturbed cancer maintained its electronegativity (Φ_1_(r, *θ, φ*) *<* 0 mV) during its growth.

For *µ*_*T*_ = 1.0, we showed results of simulations of 2D Φ_1_(r, *θ, φ*) (Fig 5) and *ρ*_*v*_(*t*) (Fig 6) space-time patterns and their respective animated movies on S2–S4 Videos in the Supporting Information. Both types of patterns were shown in Figs 5,6A (*t* = 0 days), Figs 5,6B (*t* = 5 days), Figs 5,6C (*t* = 10 days) and Figs 5,6D (*t* = 15 days) for *Y*_20_; Figs 5,6E (*t* = 0 days), Figs 5,6F (*t* = 5 days), Figs 5,6G (*t* = 10 days) and Figs 5,6H (*t* = 15 days) for *Y*_21_; Figs 5,6I (*t* = 0 days), Figs 5,6J (*t* = 5 days), Figs 5,6K (*t* = 10 days) and Figs 5,6L (*t* = 15 days) for *Y*_32_.

For *µ*_*T*_≠ 1.0 (*µ*_*T*_ *<* 1.0 and *µ*_*T*_ *>* 1.0), we present the results of simulations of 2D Φ_1_(r, *θ, φ*) (Figs 7 and 9) and *ρ*_*v*_(*t*) (Figs 8 and 10) space-time patterns and their some animated movies (see S5 Video to S7 Video in the Supporting Information). Both types of patterns were shown in Figs 7,8A (*µ*_*T*_ = 0.5 at *t* = 0 days); Figs 7,8B (*µ*_*T*_ = 0.5 at *t* = 5 days); Figs 7,8C (*µ*_*T*_ = 0.5 at *t* = 10 days); Figs 7,8D (*µ*_*T*_ = 0.5 at *t* = 15 days); Figs 7,8E (*µ*_*T*_ = 25.0 at *t* = 0 days); Figs 7,8F (*µ*_*T*_ = 25.0 at *t* = 5 days); 7,8G (*µ*_*T*_ = 25.0 at *t* = 10 days); Figs 7,8H (*µ*_*T*_ = 25.0 at *t* = 15 days); Figs 7,8I (*µ*_*T*_ = 50.0 at *t* = 0 days); Figs 7,8J (*µ*_*T*_ = 50.0 at *t* = 5 days); Figs 7,8K (*µ*_*T*_ = 50.0 at *t* = 10 days); and Figs 7,8L (*µ*_*T*_ = 50.0 at *t* = 15 days) for *Y*_20_. These patterns for *Y*_21_ were represented in Figs 9,10A (*µ*_*T*_ = 25.0 at *t* = 0 days); Figs 9,10B (*µ*_*T*_ = 25.0 at *t* = 5 days); Figs 9,10C (*µ*_*T*_ = 25.0 at *t* = 15 days); Figs 9,10D (*µ*_*T*_ = 25.0 at *t* = 15 days); Figs 9,10E (*µ*_*T*_ = 50.0 at *t* = 0 days); Figs 9,10F (*µ*_*T*_ = 50.0 at *t* = 5 days); Figs 9,10G (*µ*_*T*_ = 50.0 at *t* = 15 days); and 9,10H (*µ*_*T*_ = 50.0 at *t* = 15 days). Furthermore, simulations of both physical magnitudes were depicted in Figs 9,10I,10i (*µ*_*T*_ = 25.0 at *t* = 0 days); Figs 9,10J (*µ*_*T*_ = 25.0 at *t* = 5 days); Figs 9,10K (*µ*_*T*_ = 25.0 at *t* = 15 days); Figs 9,10L (*µ*_*T*_ = 25.0 at *t* = 15 days); 9,10M (*µ*_*T*_ = 50.0 at *t* = 0 days); Figs 9,10N (*µ*_*T*_ = 50.0 at *t* = 5 days); Figs 9,10O (*µ*_*T*_ = 50.0 at *t* = 15 days); and Figs 9,10P (*µ*_*T*_ = 50.0 at *t* = 15 days) for *Y*_32_.

Fig 11 showed how *Y*_41_ influences 2D Φ_1_(r, *θ, φ*) (Figs 11A-D) and *ρ*_*v*_(*t*) (Figs11E-H) space-time patterns. Both patterns were depicted at *t* = 0 days (Figs 11A,E), 5 days (Fig 11B,F), 10 days (Fig 11C,G) and 15 days (Fig 11D,H). Animated movies of Φ_1_(r, *θ, φ*) and *ρ*_*v*_(*t*) based on the results in Fig 11 are shown in S8 Video of the Supporting Information.

The results showed in Figs 5-11 and their respective Supporting Information S1-S9 Videos revealed that *Y*_*nm*_ and *µ*_*T*_ essentially influenced 2D Φ_1_(r, *θ, φ*) and *ρ*_*v*_(*t*) space-time patterns in each *t* during entire unperturbed cancer growth from initial value *R*_*T*_ = 5.6 mm, being marked for higher values of *µ*_*T*_, higher-order odd *Y*_*nm*_ and when *t* increased. The unperturbed cancer maintained its spherical geometry at each *t* for all values of *µ*_*T*_ and types of *Y*_*nm*_, except for *Y*_21_ and *Y*_32_ when *µ*_*T*_ = 50.0.

Although unperturbed cancer maintains its spherical shape, the concentric isolines and isosurfaces of Φ_1_(r, *θ, φ*) changed from circular/quasi-circular shapes at *t* = 0 days to non-circular shapes for *t >* 0 days that depends on the value of *µ*_*T*_ and type of *Y*_*nm*_, which was notable for the larger values of *µ*_*T*_ and *t*, as well as higher-order *Y*_*nm*_, fundamentally the most asymmetric. The simultaneous presence of intratumoral regions with negative and positive Φ_1_(r, *θ, φ*) values at *t >* 0 days was surprising and depended on the *Y*_*nm*_ type. The quantity, intensity, and intratumoral spatial distribution of these regions with negative and positive Φ_1_(r, *θ, φ*) polarities depends on the *Y*_*nm*_ type, *µ*_*T*_ value, and observation time *t*, with higher *µ*_*T*_ and *t* values corresponding to higher-order *Y*_*nm*_. These positive regions with positive Φ_1_(r, *θ, φ*) polarity were observed for *Y*_20_ and *Y*_32_ with *µ*_*T*_ = 1.0, *µ*_*T*_ = 25.0, and *µ*_*T*_ = 50.0 for *Y*_20_, *µ*_*T*_ = 50.0 for *Y*_21_, and *µ*_*T*_ = 25.0 and *µ*_*T*_ = 50.0 for *Y*_32_. The outermost isoline/isosurface of each of these positive regions had a value of Φ_1_(r, *θ, φ*) = 0 mV.

Unlike Φ_1_(r, *θ, φ*), negative *ρ*_*v*_(*t*) values were observed throughout the unperturbed cancer volume at *t* = 0 days for all *µ*_*T*_ values and *Y*_*nm*_ types, except for *Y*_32_ and *µ*_*T*_ = 25.0, and not for *t >* 0 days. Intratumoral regions with negative and positive *ρ*_*v*_(*t*) polarities were observed for all *µ*_*T*_ values, *Y*_*nm*_ type (alone or combined), and *t >* 0 days. The quantity, intensity, and spatial distribution of these negative and positive *ρ*_*v*_(*t*) regions depended on *µ*_*T*_ and *Y*_*nm*_ type, being marked for higher *µ*_*T*_ and *t* values, as well as higher-order *Y*_*nm*_.

Positive values of *ρ*_*v*_(*t*) on spherical shell of width 2*ϵ* for all *t* (Figs 4E-H), positive values of *ρ*_*v*_(*t*) on spherical shell were observed for *Y*_21_, *µ*_*T*_ = 50.0 and *t* ≥ 10 days (Figs 10G,H); *Y*_32_, *µ*_*T*_ = 25.0 and 50.0, and *t* = 15 days (Figs 10L,P). Figs 10G,H,L,P show a surprising result because *ρ*_*v*_(*t*) was positive not only in the on the spherical shell but also in most of the unperturbed cancer volume. In other words, the unperturbed cancer self-reversed its polarity from negative to positive at a certain instant *t* during its growth. In contrast, negative *ρ*_*v*_(*t*) prevailed on the spherical shell of width 2*ϵ* for the rest of the spatiotemporal distributions of this physical quantity for all *t* (Figs 6,8A-L; 10A-D and 11E-H). Furthermore, negative *ρ*_*v*_(*t*) prevailed on spherical shell of width 2*ϵ* for *Y*_21_, *µ*_*T*_ = 50.0 and *t <* 10 days (Figs 10G,E,F); *Y*_32_, *µ*_*T*_ = 25.0 and 50.0, and *t <* 15 days (Figs 10I-K,M-O).

Figs 4-11 show that 2D Φ_1_(r, *θ, φ*) and *ρ*_*v*_(*t*) spatiotemporal patterns were more heterogeneous for higher values of *µ*_*T*_ and more asymmetrical *Y*_*nm*_ (higher-order odd *Y*_*nm*_), which was marked as *t* increased. Nevertheless, the morphologies and symmetry of both patterns were conserved during unperturbed cancer growth (*t >* 0 days). This result is valid when one (Figs 5-11) or several (Fig 4) *Y*_*nm*_ were included in our approach.

The results show in Figs 4, 6, 8, 10, and 11 evidenced that negative (red color) and positive (blue color) values of *ρ*_*v*_(*t*) corresponded to inhibitor (blue color) and activator (red color) mechanisms in Turing patterns for each *Y*_*nm*_ (Fig 2), respectively.

Fig 12 shows the global time behaviors of *R*_*T*_ (Figs 12A,D,G) and ||*ρ*_*v*_(*t*)|| norm (Figs 12B,E,H), as well as the phase diagram of ||*ρ*_*v*_(*t*)|| versus *R*_*T*_ (Figs 12C,F,I) for *Y*_20_, *Y*_21_, *Y*_32_ and *µ*_*T*_ = 0.5, 1.0, 25.0 and 50.0. Fig 13 represents the global time behavior of ||*ρ*_*vs*_(*t*)|| norm (Figs 13A,C,E) and the phase diagram of ||*ρ*_*vs*_(*t*)|| versus *R*_*T*_ (Figs 13B,D,F) for *Y*_20_, *Y*_21_, *Y*_32_ and *µ*_*T*_ = 1.0, 25.0 and 50.0. Several findings were revealed in these two figures First, *R*_*T*_ increases with *t* for all *µ*_*T*_ and *Y*_*nm*_, being marked for *µ*_*T*_ = 50.0 and *Y*_32_. Second, ||*ρ*_*v*_(*t*)|| and ||*ρ*_*vs*_(*t*)|| norms increases with *t* or *R*_*T*_ and depended on *µ*_*T*_ and *Y*_*nm*_ type. The highest values of these two norms were obtained for *µ*_*T*_ = 25.0 and *Y*_20_ (Figs 12B,C and 13A,B), *µ*_*T*_ = 50.0 and *Y*_21_ (Figs 12E,F and 13C,D), *µ*_*T*_ = 1.0 and *Y*_32_ (Figs 12H,I and 13E,F). Third, Figs 12A-F and 13A-D showed that higher value of || *ρ*_*v*_(*t*) || and || *ρ*_*vs*_(*t*) || norms correspond to faster unperturbed TGK, whereas slower unperturbed TGK correspond to lower values of these two norms, as expected. In contrast, *R*_*T*_ grew very rapidly over *t* and lower values of || *ρ*_*v*_(*t*) || and || *ρ*_*vs*_(*t*) || norms were observed for *Y*_32_ and *µ*_*T*_ = 50.0, a surprising result (Figs 12G,H,I and 13E,F).

**Fig 12.**
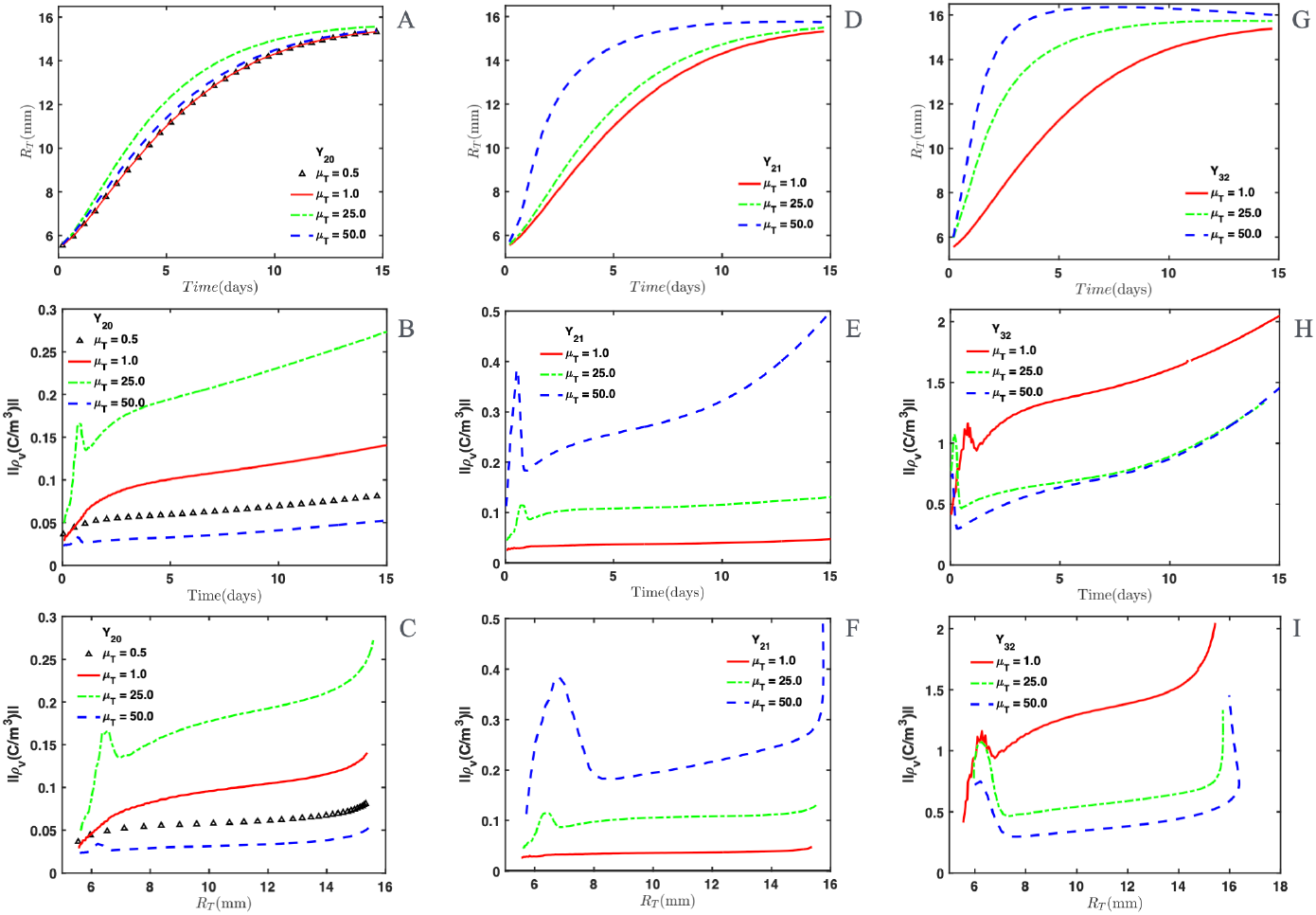
Surface standard metrics. Norm ||*ρ*_*v*_(*t*)|| for each value of mass *µ*_*T*_ and three spherical harmonics (*Y*_20_, *Y*_21_ and *Y*_32_). (A) Plot of *R*_*T*_ versus *t* for *Y*_20_. (B) Plot of ||*ρ*_*v*_(*t*)|| versus *t* for *Y*_20_. (C) Plot of ||*ρ*_*v*_(*t*)|| versus *R*_*T*_ for *Y*_20_. (D) Plot of *R*_*T*_ versus *t* for *Y*_21_. (E) Plot of ||*ρ*_*v*_(*t*)|| versus *t* for *Y*_21_. (F) Plot of ||*ρ*_*v*_(*t*)|| versus *R*_*T*_ for *Y*_21_. (G) Plot of *R*_*T*_ versus *t* for *Y*_32_. (H) Plot of ||*ρ*_*v*_(*t*)|| versus *t* for *Y*_32_. (I) Plot of ||*ρ*_*v*_(*t*)|| versus *R*_*T*_ for *Y*_32_.

**Fig 13.**
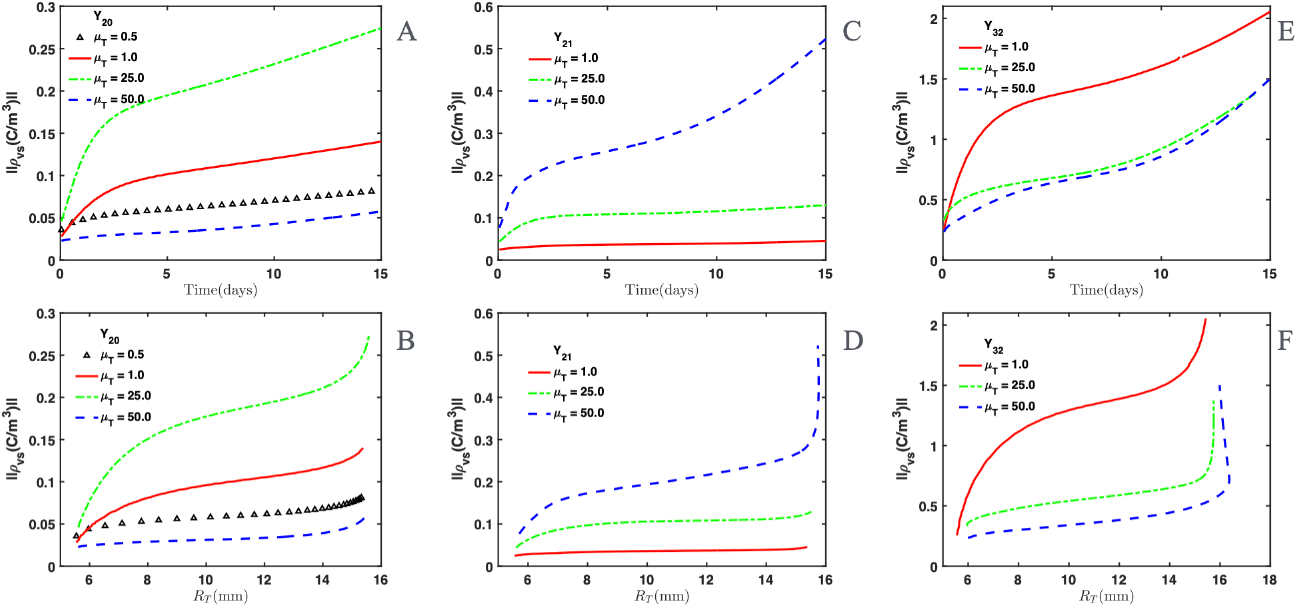
Volume standard metrics. Norm ||*ρ*_*vs*_(*t*)|| for each value of mass *µ*_*T*_ and three spherical harmonics (*Y*_20_, *Y*_21_ and *Y*_32_). (A) Plot of ||*ρ*_*vs*_(*t*)|| versus *t* for *Y*_20_. (B) Plot of ||*ρ*_*vs*_(*t*)|| versus *R*_*T*_ for *Y*_20_. (C) Plot of ||*ρ*_*vs*_(*t*)|| versus *t* for *Y*_21_. (D) Plot of ||*ρ*_*vs*_(*t*)|| versus *R*_*T*_ for *Y*_21_. (E) Plot of ||*ρ*_*vs*_(*t*)|| versus *t* for *Y*_32_. (F) Plot of ||*ρ*_*vs*_(*t*)|| versus *R*_*T*_ for *Y*_32_.

A fifth finding revealed in Figs 12 and 13 is the peak (Figs 12B,C,E,F,H,I) or slope change (Figs 13A-F) of ||*ρ*_*v*_(*t*)|| and ||*ρ*_*vs*_(*t*)|| norms observed in the early times of *t*.

Another finding shown in these two figures was the rapid (Figs 12B,E,H and 13A,C,E) or abrupt (Figs 12C,F,I and 13B,D,F) slope change of ||*ρ*_*v*_(*t*)|| and ||*ρ*_*vs*_(*t*)|| norms, although *R*_*T*_ tends to an asymptotic value for long *t*.

## Discussion

This study proposes a mechanical-electrical formalism that allows us to theoretically understand how *Y*_*nm*_ (single or combined) and *µ*_*T*_ affect 2D Φ_1_(r, *θ, φ*), *ρ*_*v*_(*t*) and *σ*_12_ (at Σ) space-time distributions, unperturbed TGK (*R*_*T*_ versus *t*), and values of ||*ρ*_*v*_(*t*)|| and || *ρ*_*vs*_(*t*) || for each time instant *t*. Nevertheless, it has three fundamental limitations. First, our theoretical results have not been corroborated in the experiment. Second, the spherical shape of unperturbed cancer because it generally adopts a non-spherical shape, as observed in *in vitro* [71], *ex vivo* [72], preclinical [7, 8, 73–75] and clinical [23, 69, 76] studies. Despite this second limitation, spherical cancer is widely used in experimental (*in vitro, ex vivo* and *in vivo*) [14, 77–81] and theoretical [11, 13, 82, 83] studies. Third, Φ_0_ and Φ_*s*_ are fixed in our simulations during entire unperturbed TGK. Although this third limitation has not been corroborated or refuted in experimental and theoretical studies, we expect Φ_0_ and Φ_*s*_ must change over time during unperturbed cancer growth; therefore, more complex Φ_1_(r, *θ, φ*), *ρ*_*v*_(*t*) and *σ*_12_ spatiotemporal patterns.

Despite the three limitations mentioned above, the good correspondence between the curves showed in *σ*_12_ versus *R*_*T*_ and *dσ*_12_*/dt* versus *σ*_12_ plots for the same value of *CDR* and *CDR*_*PF*_ confirm theoretical results reported in [13] and the correct introduction of *Y*_*nm*_ and *µ*_*T*_ in our physical-mathematical approach for the analysis of Φ_1_(r, *θ, φ*), *ρ*_*v*_(*t*), *σ*_12_, ||*ρ*_*v*_(*t*)|| and ||*ρ*_*vs*_(*t*)||, as showed in Fig 4. Furthermore, our simulations of Φ_1_(r, *θ, φ*) mimic experimental results reported by Miklavčič et al. [26] when *R*_*T*_ = 5.6 mm at *t* = 0 days for all *µ*_*T*_ and *Y*_*nm*_. This is argued by the following reasons: 1) 2D Φ_1_(r, *θ, φ*) is more negative in unperturbed cancer central region and less negative towards its contour/surface Σ. 2) Concentric quasi-circular/quasi-spherical 2D Φ_1_(r, *θ, φ*) isolines/isosurfaces may explain the experimental values of this physical magnitude along axial and radial directions documented in [26]. 3) Negative values of Φ_1_(r, *θ, φ*) and *ρ*_*v*_(*t*) throughout the entire volume (interior and contour/surface) of unperturbed cancer at *t* = 0 days confirm its electronegativity (Φ_1_(r, *θ, φ*) *<* 0, *ρ*_*v*_(*t*) *<* 0 and *σ*_12_ *<* 0), as in experiment [14, 26, 67, 84] and theory [13].

Although Φ_1_(r, *θ, φ*), *ρ*_*v*_(*t*) and *σ*_12_ values have not been experimentally measured, theoretical values of *σ*_12_ for Sa-37 fibrosarcoma unperturbed cancer in range of -0.0080 to -0.0005 *C/cm*^2^, dependent on Φ_0_ − Φ_*s*_ and *CDR*/*CDR*_*PF*_ values, are consistent with those reported experimentally in fibroblasts (*σ*_12_ ≈ − 0.0125 *C/cm*^2^), MCF-7 breast cancer cells (*σ*_12_ ≈ − 0.0200 *C/cm*^2^), and MDA-MB-231 breast cancer cells (*σ*_12_ ≈ − 0.0175 *C/cm*^2^) for pH values in the range of 6-7 [85]. It is important to note that pH range measured in unperturbed cancer corresponds to 6–7 [86].

Our simulations reveal that the range of *σ*_12_ for Sa-37 fibrosarcoma unperturbed cancer is smaller than that documented in [85] for fibroblasts, MCF-7 breast cancer cells and MDA-MB-231 breast cancer cells. This may be explained because Sa-37 fibrosarcoma cancer cells are more aggressive, have lower transmembrane potential and higher electrical conductivity in comparison with fibroblasts and two breast cancer cell lines. The later is consistent with previous studies [7, 11, 13, 15–17, 23, 26, 73, 84, 87].

The correct introduction of *Y*_*nm*_ and *µ*_*T*_ in our theoretical approach allows us to reveal new aspects of unperturbed cancer bioelectricity from a theoretical point of view and its direct connection with unperturbed TGK and cancer biology. All these bioelectrical and biological aspects in unperturbed cancer are markedly influenced by *Y*_*nm*_ and *µ*_*T*_, unprecedented in the literature; hence, the following merits of this study that are closely related to each other, which are listed below in order of importance.

The first merit lies in the close connection among dynamic physicochemical processes (Turing patterns obtained with BVMA model for different *δ* values); anomalous bioelectricity (*η*_1_, *ε*_1_, *η*_2_, *ε*_2_, Φ_1_(r, *θ, φ*), *ρ*_*v*_(*t*) and *σ*_12_); biomechanical (*ϕ, B*_*ϕ*_ and ℳ [Φ_1_]); and entire TGK (*G*(*t*) and *R*_*T*_ versus time) in unperturbed cancer. Second merit, entire unperturbed TGK are essentially self-regulated by two closely interconnected parameters: degree of *Y*_*nm*_ asymmetry and *µ*_*T*_ value concerning 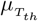 (threshold value of *µ*_*T*_ ). Third merit, inhibition and activation unperturbed cancer growth mechanisms (Turing patterns) are related to self-regulated dynamic regions with negative and positive Φ_1_(r, *θ, φ*), *ρ*_*v*_(*t*) and *σ*_12_ polarities, which depend markedly on *Y*_*nm*_ (alone or in combination), *µ*_*T*_ and *t*. Fourth merit, anomalous bioelectricity of unperturbed cancer depends on its dynamic mechanoelectrical nature (*η*_1_, *ε*_1_, *η*_2_, *ε*_2_, Φ_1_(r, *θ, φ*), *G*(*t*), *ρ*_*v*_(*t*), *σ*_12_, *ϕ, B*_*ϕ*_, degree of asymmetry of *Y*_*nm*_, *µ*_*T*_ value with respect to 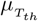, and *t*), and not only on intratumoral faradic (associated to electrons) and ionic (linked to ions) currents, as report in [15].

The physical-chemical-biological-growth kinetic processes are inextricably linked in unperturbed cancer and cannot be analyzed separately, as the close connection among abnormal bioelectricity, mechanical-morphological and hallmarks, a matter that agree with previous studies [7, 8, 11, 13, 15]. This assertion may be explained by these two aspects derived from our theoretical approach. First, unperturbed TGK (*R*_*T*_ versus *t*), geometry (spherical or non-spherical) and biological heterogeneity of unperturbed cancer are influenced by 2D Φ_1_(r, *θ, φ*), *ρ*_*v*_(*t*), *σ*_12_ and Turing spatiotemporal patterns, as well as global values of *ρ*_*v*_(*t*) (||*ρ*_*v*_(*t*)||/||*ρ*_*vs*_(*t*)||), which in turn depend on whether *Y*_*nm*_ is symmetric or asymmetric and *µ*_*T*_ value regarding 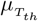. Second, temporal behaviors of *R*_*T*_ are similar for symmetrical *Y*_*nm*_ (e.g., *Y*_20_) and all *µ*_*T*_ value, but not for asymmetrical *Y*_*nm*_, as *Y*_21_, *Y*_32_ and *Y*_41_ (see Supporting Information). Consequently, unperturbed cancer is mechanical and electrical in nature since primary physicochemical changes bring about biological changes, which in turn lead to clinical changes, in agreement with [7, 8, 11, 13, 23]. Third, the mechanical-electrical connection proposed in the energy model confirms that charge density-induced unperturbed cancer surface generation is essential to minimize energy costs during its growth (due to the term ∇*ϕ* · ∇*B*_*ϕ*_). Furthermore, the internal self-regulation of *ρ*_*v*_(*t*), determined by a biochemical pattern (from the BVAM model), generates directed growth where energy costs are lower. Likewise, areas with lower mean curvature (large *B*_*ϕ*_) promote electric reorganization to favor unperturbed growth. Ultimately, the biochemical-electrical behavior exhibits self-regulation, promoting unperturbed cancer growth in areas of lower energy cost and higher cellular activity, whereas simultaneously sacrificing growth to increase bielectricity in the periphery and interior of unperturbed cancer. Of note, the time-neutral reorganization of *ρ*_*v*_(*t*) corresponds to an inhibitory process that on average remains zero, but as a consequence generates spatiotemporal modifications in unperturbed cancer to mechanically and electrically compensate its evolution.

A contradiction between the time increase of *R*_*T*_ and the decrease of ||*ρ*_*v*_(*t*)||/||*ρ*_*vs*_(*t*)|| in *t* is observed for 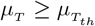, *t >* 5 days and both symmetrical and asymmetrical *Y*_*nm*_, being marked for *Y*_*nm*_ asymmetric, unexpected and surprising theoretical result, which would correspond to highly undifferentiated (or highly aggressive) unperturbed cancer in the experimental order. This contradiction would imply that unperturbed cancer would self-regulate its fast growth by two possibilities depending on whether *Y*_*nm*_ is symmetric or asymmetric as *R*_*T*_ increases over time *t*. For symmetrical *Y*_*nm*_, negative values of Φ_1_(r, *θ, φ*) and *ρ*_*v*_(*t*) would decrease continuously to slightly negative values close to zero due to the overlap of intratumoral regions with slightly negative and positive values of these two physical magnitudes in regions of the spherical cap and surface of unperturbed cancer. This would allow the entry of cellular and humoral elements of the immune system into interior of it, probably through the positive regions of Φ_1_(r, *θ, φ*) and *ρ*_*v*_(*t*). Nevertheless, for asymmetrical *Y*_*nm*_, negative values of Φ_1_(r, *θ, φ*) and *ρ*_*v*_(*t*) would change drastically to positive, being marked for more asymmetric *Y*_*nm*_. This would mean that in the same unperturbed TGK, the cancer would change from very undifferentiated (*t* ≤ 5 days) to well-differentiated, behave like the surrounding healthy tissue positively polarized from which it originated, or most of its volume is in a quiescent or dormant permanent state; therefore, its cells do not enter the cell cycle (*t >* 5 days). For both symmetrical and asymmetrical *Y*_*nm*_, unperturbed cancer would self-destruct or grow slowly over time time during the second stage of TGK and would take a long time to reach the steady state (third stage of unperturbed TGK) [52], in contrast with experiment [7, 52, 73, 74, 88, 89]. This contradiction is not observed when 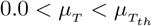 for all *t* and *Y*_*nm*_ type; therefore, we focus in the later case.

The results of this study suggest that the influence local changes in Φ_1_(r, *θ, φ*), *ρ*_*v*_(*t*), *σ*_12_ and ||*ρ*_*v*_(*t*)||/||*ρ*_*vs*_(*t*)|| do not necessarily lead to changes in the global behavior of unperturbed TGK, as evidenced in *R*_*T*_ versus *t* plot for *Y*_20_ and any *µ*_*T*_ value, not so for odd asymmetric *Y*_*nm*_, as shown in *R*_*T*_ versus *t* plot for *Y*_21_, *Y*_32_ and *Y*_41_ and larger values of 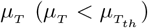. This confirm that unperturbed cancer may experience local changes rather than global changes due to its complexity and the interactions that occur at different scales, in agreement with [90, 91]. Furthermore, the diversity of *R*_*T*_ behaviors in terms of *Y*_*nm*_ and *µ*_*T*_ value, provided that 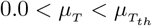, may suggest that these two parameters are in correspondence with the degree of cancer differentiation. Differentiated or well differentiated unperturbed cancer may correspond to that with symmetrical *Y*_*nm*_ and any value of *µ*_*T*_ . Moderately differentiated unperturbed cancer may be that with odd *Y*_*nm*_ and *µ*_*T*_ ≤ 1.0. Undifferentiated unperturbed cancer may be consonant with odd *Y*_*nm*_ of higher order *n* and 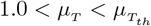.

Since negative values of *ρ*_*v*_(*t*) contribute more than positive values to ||*ρ*_*v*_(*t*) || / ||*ρ*_*vs*_(*t*) ||, the increase and prevalence of regions with negative *ρ*_*v*_(*t*) values throughout unperturbed cancer volume leads to its faster TGK and vice versa, as observed when 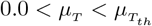 for asymmetric *Y*_*nm*_ and higher *µ*_*T*_ value. This is verified because *R*_*T*_ growth rate increases with the increase in ||*ρ*_*v*_(*t*)||/||*ρ*_*vs*_(*t*)|| depending on *µ*_*T*_ value and degree of *Y*_*nm*_ asymmetry. This diversity of time behaviors for *R*_*T*_ may mimic those observed for well-differentiated, moderately differentiated, and undifferentiated unperturbed cancer in clinics [69, 89]. We think that dynamic self-regulations of intratumoral regions with negative and positive polarities of Φ_1_(r, *θ, φ*), *ρ*_*v*_(*t*) and *σ*_12_, as well as temporal behaviors of ||*ρ*_*v*_(*t*)||/||*ρ*_*vs*_(*t*)|| and Turing spatiotemporal patterns not only influence *G*(*t*) and *R*_*T*_ versus *t*, but may also have relevant implications for unperturbed cancer biophysics, such as: 1) maximum survival (a key characteristic of biological tissues and organisms, as open and nonlinear systems) [7, 52]. 2) Self-maintenance, self-organization, and self-regulation of all its vital biophysical-chemical processes (e.g., bioelectricity, central or non-central endogenous intratumoral necrosis, intratumoral electrical heterogeneity, metastasis) and in the symmetry conservation over time of Φ_1_(r, *θ, φ*), *ρ*_*v*_(*t*), *σ*_12_ and Turing spatial patterns throughout entire unperturbed TGK. 3) Protection of unperturbed cancer against attack by external agents (e.g., immune system and anticancer therapies).

Endogenous intratumoral necrosis regions (central and non-central) in unperturbed cancer [14, 69, 88] has been explained by different mechanisms that are not independent and must be interconnected, such as: 1) endogenous blood vessels do not reach these regions, mainly central region; therefore, neither do nutrients and oxygen [5, 69, 89]. 2) Intense electric fields (due to more electronegative electrical potentials) break weak bonds between cancer cells [7, 68], as report in [13]. 3) Cancer cells diffuse from their central region to the periphery [11, 14, 41], which may be consistent with changes in the Turing pattern over time. Furthermore, central or non-central necrosis endogenous intratumoral necrosis are indicators of fast-growing and aggressive of unperturbed cancer, greater metastatic spread of cancer cells, and poor prognosis for cancer organism, being marked for non-central necrosis [5, 69]. These two forms of endogenous intratumoral necrosis presentation within unperturbed cancer may be corroborated from our simulations when their results are compared when those reported experimentally and theoretically by Serŝa *e*t. al [14] for symmetric Sa-1 sarcoma growing in male A/J mice (Figs 2a,b) and asymmetric Sa-1 sarcoma growing on male A/J mice (Figs 2a,b) and T50/80 mammary carcinoma growing on immunodeficient mice (Figs 3a,e). The necrotic and active intratumoral regions are represented by dark (low signal by MRI and high current density by CDI) and bright (high signal by MRI and low current density by CDI) colors, respectively [14].

In Fig 14 central necrotic and active (in spherical cap near cancer surface) endogenous intratumoral regions in symmetric Sa-1 sarcoma may be due to the high (red color in *ρ*_*v*_(*t*) spatiotemporal pattern) and low (blue color in *ρ*_*v*_(*t*) spatiotemporal pattern) concentrations of negative charge carriers in the central region and spherical cap (including the cancer surface), respectively. Central necrotic region is consistent with inhibitor mechanism (blue color in Turing pattern) and high unperturbed cancer density (red color in [11, 41]), whereas active region correspond with activator mechanism (red color in Turing pattern) and low unperturbed cancer density (blue color in [11, 41]). These space-time patterns may explain similarity among the temporal behaviors of *R*_*T*_ for symmetrical *Y*_*nm*_ (e.g., *Y*_20_) and all *µ*_*T*_ (see animated movie on S9 Video in Supporting Information). Furthermore, red and blue colors in *ρ*_*v*_(*t*) spatiotemporal pattern corresponded with regions of high and low electrical conductivities in 3D electrical impedance tomography imaging, respectively [92–94].

**Fig 14.**
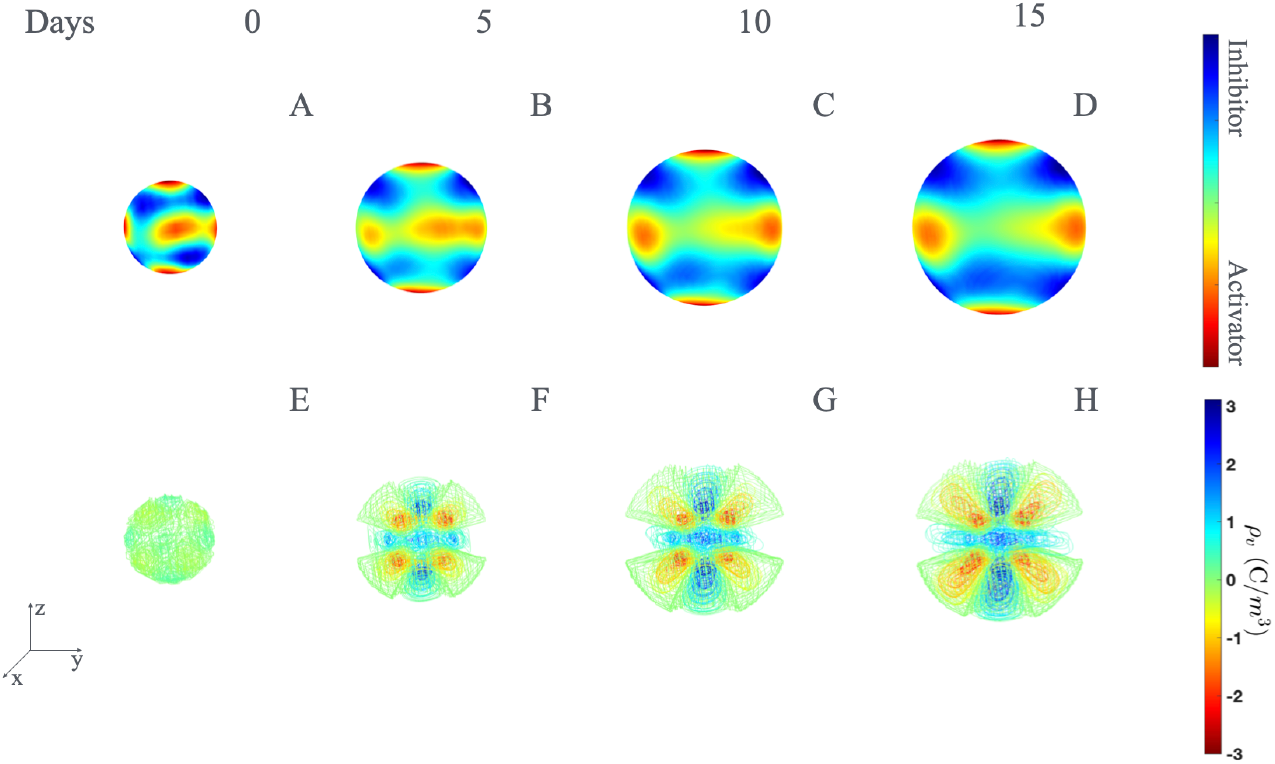
Simulations of BVAM model for stripe patterns. Turing patterns in 3D (A-D) and charge density *ρ*_*v*_(*t*) spatiotemporal patterns (E-H) obtained with Eqs (18) and (28). These patterns were showed at *t* = 0 days (A,E), *t* = 5 days (B,F), *t* = 10 days (C,G) and *t* = 15 days (D,H) for *µ*_*T*_ = 1.0, *c* = 0.00 and *δ* = 1.55.

Non-central necrosis (a focus or multiple foci of necrotic tissue outside the central region) and active endogenous intratumoral necrosis in asymmetric Sa-1 sarcoma and T50/80 mammary carcinoma could be explained from the number of regions that contain negatively (*ρ*_*v*_(*t*) *>* 0) and positively (*ρ*_*v*_(*t*) *<* 0) charged carriers, which are distributed throughout entire unperturbed cancer volume, depending on degree of *Y*_*nm*_ asymmetry and *µ*_*T*_ value, being marked for *Y*_*nm*_ more asymmetrical. The later is consistent with Turing spatiotemporal pattern. These space-time patterns explain differences among *R*_*T*_ behaviors for different *µ*_*T*_ values and degree of *Y*_*nm*_ asymmetry, being marked for *Y*_*nm*_ more asymmetrical, which induce fastest unperturbed TGK, as observed for asymmetric unperturbed cancer in preclinical studies [7, 8, 52, 88].

On the other hand, the dark and bright stripes and gray levels (combinations of black and white) observed by Serŝa *e*t. al [14] for symmetric or asymmetric unperturbed cancer are theoretically mimicked by color stripes observed in Φ_1_(r, *θ, φ*), *ρ*_*v*_(*t*), *σ*_12_ and Turing spatiotemporal patterns (color tones between red and blue), as shown in Fig 14 for space-time patterns of Turing (Figs 14A-D) and *ρ*_*v*_(*t*) (Figs 14E-H). These stripes are also consistent with the multiple density layers of symmetric (color tones between red and blue) and asymmetric (different tones of blue color) unperturbed cancer documented in [11, 41]. Consequently, the color intensity in each of these stripes may be related to the *µ*_*T*_ value.

Dynamic self-regulations of intratumoral regions with negative and positive of 2D Φ_1_(r, *θ, φ*), *ρ*_*v*_(*t*) and *σ*_12_ values, as well as intratumoral density [11, 41], necrosis (central or non-central) [14, 69, 88] and Turing spatiotemporal patterns may explain intratumoral electrical heterogeneity and anisotropy, which are marked for odd *Y*_*nm*_ more asymmetric, higher *µ*_*T*_ value and longer *t*. Both intratumoral electrical heterogeneity and anisotropy may act as a heterogeneous intratumoral electrical shield and suppose that cancer cells with different electrical properties (e.g., electrical conductivity and permittivity) and biological characteristics (e.g., shape, phenotype and genotype) concentrate in multiple concentric layers, as report in [13], or in several discrete electric sub-shields distributed conveniently throughout unperturbed cancer depending markedly on the degree of asymmetry of *Y*_*nm*_.

Both intratumoral electrical heterogeneity and anisotropy lead to intratumoral biological heterogeneity and anisotropy of unperturbed cancer [7, 8, 13, 22, 24, 38, 95–98], which constitute cancer hallmarks [5, 6], in agreement with our simulations: more asymmetric odd *Y*_*nm*_ (greater heterogeneity and electrical anisotropy in unperturbed cancer) results in faster unperturbed TGK (greater heterogeneity and electrical anisotropy in unperturbed cancer). This intratumoral biological heterogeneity may result in intrinsic anisotropic patterns in unperturbed cancer, as report experimentally [99–101] and theoretically [11, 41] in both spherical and non-spherical tumors. The biological heterogeneity and anisotropy in spherical unperturbed cancer could be related to anisotropic diffusive flow [102], intratumoral stresses [95], matrix stiffness [103], drug resistance [104], among others.

Deformation of 2D Φ_1_(r, *θ, φ*) isolines/isosurfaces, *ρ*_*v*_(*t*) isosurfaces and Turing pattern may explain the loss of unperturbed cancer sphericity observed for asymmetric *Y*_*nm*_, higher *µ*_*T*_ value and 0 *< t <* 5 days, in agreement with the phase transition between avascular and vascular phases of unperturbed TGK, in which cancer shape changes from spherical to ellipsoidal, as document in Ehrlich and Sa-37 fibrosarcoma tumors [7, 8, 52, 105], F3II mammary carcinoma [28]) and theoretical simulations [11]. We believe that this phase transition is physical in nature, probably of a mechanical-electrical type, because it may be related to local Avrami coefficient (*n*_*loc*_) sharp peak [7]; cancer density [11]; hard phase transition type, as “first order” by means of a supercritical Andronov–Hopf bifurcation, emergence of limit cycle, and a cascade of bifurcations, as that of saddle-foci (Shilnikov [106]); and *ρ*_*v*_(*t*) peak, whose height and width depend on *µ*_*T*_ and *Y*_*nm*_ type, as we show in ||*ρ*_*v*_(*t*)|| versus *t* and ||*ρ*_*v*_(*t*)|| versus *R*_*T*_ plots.

The stationary stage for long *t* or third stage of TGK [52], verified in Eq (27), has been explained from the balance between the rates of cancer cell production and loss [52, 69, 89]; nutrient and oxygen limitation, mechanical pressure and limited space, accumulation of metabolic waste, and host immune response [58, 107, 108]. Nevertheless, our simulations suggest that this stationary stage is self-regulated by the unperturbed cancer itself from existence of intratumoral *ρ*_*v*_(*t*) negative and positive regions, which could be related to cell loss processes (e.g., apoptosis, necrosis, exfoliation and metastasis [69, 89]) and mechanism that activates cell duplication to replenish the number of lost cancer cells, respectively. We believe that cell loss processes are related to inhibitor mechanism in Turing pattern, whereas activating processes of the cell cycle may be related to activating mechanisms in Turing pattern.

Despite the electrical and biological heterogeneities and anisotropies in unperturbed cancer, it dynamically self-regulates electronegativity and electropositivity of different intratumoral regions to self-compensate/self-balance global electronegativity throughout its volume (*ρ*_*v*_(*t*) *<* 0) and self-maintain symmetry of 2D Φ_1_(r, *θ, φ*), *ρ*_*v*_(*t*), *σ*_12_ and Turing spatial patterns over time, depending on the degree of asymmetry of *Y*_*nm*_. We believe that electronegativity and symmetry dynamically self-regulated in unperturbed cancer are vital for its maximum survival, self-preservation as an autonomous system in interaction with host, rapid growth and contour/surface widening self-regulation, self-preserve *σ*_12_ *<* 0, self-similarity, morphological and structural self-organization, self-adaptation, fractality processes, metastasis, and self-protection against external agents, in agreement with other studies. [7, 11, 75, 90, 91, 109–115]. High symmetry in unperturbed cancer has been related to its coherence (all its elements/components are interconnected and function in an unified manner) [75, 115].

If electronegativity is not self-regulated by electropositivity regions, unperturbed cancer would be strongly electronegative throughout its entire volume, leading to high instabilities within it; therefore, entire cancer would self-destruct, in agreement with [13], who suggest that these instabilities may be due to release of hydrogen ions (*H*^+^) from unperturbed cancer interior into its environment, whose pH becomes more acidic, an essential aspect for protection and growth of unperturbed cancer. We do not discard that this loss of *H*^+^ ions may be self-compensated by the accumulation of other positive charge carriers, such as: sodium 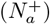, potassium (*K*^+^), magnesium 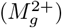, calcium 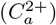, iron 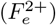, copper 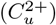, zinc 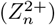, and manganese (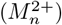 ions, which have been associated with the initiation and progression of gastric cancer [116]. We believe that intratumoral regions that contain these ions may correspond with those with positive *ρ*_*v*_(*t*) values (red color) and activating mechanism in the Turing pattern (blue color).

Simulations of Φ_1_(r, *θ, φ*), *ρ*_*v*_(*t*), *σ*_12_ and Turing spatiotemporal patterns may suggest that different negatively charged intratumoral regions, depending on the degree of *Y*_*nm*_ asymmetry and *µ*_*T*_ value, are the most likely sites for the migration of cancer cell micro-clones (or micro-clusters), instead of cell-to-cell migration, from unperturbed cancer interior into nearby and distant normal tissues, process named metastasis, in agreement with other authors [117, 118]. These negatively charged intratumoral regions become unstable at the moment that these micro-clones detach from primary unperturbed cancer. We believe that these instabilities may be compensated by intratumoral regions with *ρ*_*v*_(*t*) *>* 0, which, in turn, may serve as protective barriers for unperturbed cancer against external agents (e.g., organism as a whole and action of anticancer therapies) during its growth. The metastasis process has been explained from epithelial-mesenchymal transition [117–119]. Nevertheless, we propose that electronegativity in intratumoral region near unperturbed cancer surface (spherical cap and surface) is self-regulated in space-time during its growth according to blood (complex dielectric tissue) electrical charge density ((*ρ*_*v*−*b*_(*t*))). The *ρ*_*v*−*b*_(*t*) is not constant over time because it depends on how body temperature, hematocrit, and the concentration of electrolytes in the plasma vary in time.

We think that *ρ*_*v*_(*t*) must satisfy the following condition *ρ*_*v*_(*t*) ≊ *ρ*_*v*−*b*_(*t*) or *ρ*_*v*_(*t*) ≥ *ρ*_*v*−*b*_(*t*) to ensure that cells that migrated from unperturbed cancer enter the bloodstream through new blood vessels formed from pre-existing vessels (process known as angiogenesis, which is another hallmark of cancer [5]); otherwise, this process may not be favored. This is postulated based on the fact that blood is an electrical conductor due to the presence of ions in its plasma (chloride ions (*Cl*^−^), 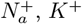 and 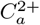), primarily from dissolved salts such as *N*_*a*_*Cl*). Although we do not know which of these two possible conditions occurs in the experiment, the first condition may be more favorable due to the potential electrical coupling between unperturbed cancer and the blood contained in newly formed blood vessels. A further study will be required to determine which of these two conditions is more favorable.

We do not rule out that number of newly formed blood vessels in unperturbed cancer depends on the number of electronegative regions near to Σ, given by odd *Y*_*nm*_ asymmetry. This could be argued because the most aggressive unperturbed cancers are usually the most angiogenic and *R*_*T*_ grows faster for asymmetric odd-*Y*_*nm*_ and higher *µ*_*T*_ value. These two later aspects suggest that spatiotemporal pattern adopted by unperturbed cancer vascular network during its growth also be related to the degree of *Y*_*nm*_ asymmetry and *µ*_*T*_ value 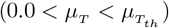. The connection between these aspects may be established from the results reported in [8], who report that unperturbed cancer surface is totally disconnected (*d*_*f*_ *<* 1), which implies the existence of pores/tunnels that are covered by newly formed blood vessels in electronegative regions near to Σ, depending on the degree of asymmetry of odd *Y*_*nm*_. Furthermore, all these surface aspects are related to the widening of the spherical cap observed for *Y*_*nm*_ with greater asymmetry, as *Y*_32_ and *Y*_41_. The later may be explained because the increase of order *n* of *Y*_*nm*_ brings about that *ρ*_*v*_(*t*) and *σ*_12_ oscillate more rapidly over the tumor surface (Σ), resulting in a more complex shape due to higher asymmetry that could be related to greater deformation of the tumor surface, a greater number of spicules formed in this surface, and widening of the tumor border, as observed in clinics for aggressiveness unperturbed cancer [23, 69]. This widening of the spherical cap confirms that the electrically and biologically active part of unperturbed cancer is its periphery, consistent with experiment [14].

Since *µ*_*T*_ is related to the aggregation of mass to unperturbed cancer, we suggest that *µ*_*T*_ and *d*_*f*_ [8] are connected. This is explained because *µ*_*T*_ and *α* are closely connected by means of the order parameter *ϕ*, and *α* is related to *d*_*f*_ . Consequently, we believe that 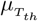 must be associated with a threshold of *d*_*f*_, called 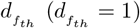, which defines the existence (*d*_*f*_ *<* 1 and 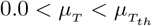) or non-existence (*d*_*f*_ ≥ 1 and 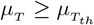 ) of unperturbed cancer. A further study is required to demonstrate the association of *µ*_*T*_ with *d*_*f*_ . We do not rule out that *d*_*f*_ is also related to *Y*_*nm*_. The latter is explained because *d*_*f*_ serves as a quantitative measure of the complexity and irregularity of unperturbed cancer contour/surface, which in turn characterizes its asymmetry and malignancy potential [7, 8, 14, 28] that are related to degree of asymmetry of *Y*_*nm*_ and *µ*_*T*_ value.

The extravasation of these micro-clones of cancer cells may be favored by blood electronegativity, which can be explained by the fact that most of its components (e.g., red blood cells, white blood cells, platelets, and plasma proteins) have a net negative surface charge [120]. Negative charges on the red blood cell surface (by presence of the carboxyl group of sialic acids in the cell membrane) induces a force that acts on the negatively charged cancer cells, causing them to travel through the bloodstream to one or more host sites that have electrical charge density (*ρ*_*v*−*h*_(*t*)) equal to or similar to those of the circulating micro-clones (*ρ*_*v*−*cc*_(*t*)), but of opposite polarity, that is, *ρ*_*v*−*h*_(*t*) = − *ρ*_*v*−*cc*_(*t*). From our simulations, we do not rule out that *ρ*_*v*−*cc*_(*t*) ≈ *ρ*_*v*_(*t*).

All this suggests that both positive and negative intratumoral regions in unperturbed cancer are related to new concepts of symmetry and symmetry breaking in cancer reported in [110]. This argued because negative intratumoral regions tend to break symmetry and cause instabilities on unperturbed cancer surface during the release of micro-clones of cancer cells, aspect relevant to its homeostasis loss, origin, growth, invasion, spread, and resistance to immune system and anticancer therapies, in agreement with Frost et al. [110]. The symmetry breaking that undergo unperturbed cancer corresponds to instability close to a first-order (Andronov-Hopf bifurcation) [106, 119] or second-order critical point (obtained by real order parameter equations in the form of Swift–Hohenberg type, which is applied to elastic materials) [91, 121]. The later is expected because unperturbed cancer is a type of elastic material [7, 8, 11, 37, 122]. We think that symmetry breaking could be self-compensating by positive intratumoral regions to maintain unperturbed cancer symmetry over time, in agreement with publish researcher [90, 111, 115].

In the experiment, the conservation of unperturbed cancer symmetry during its growth, for the same type of *Y*_*nm*_ and fixed *µ*_*T*_ value, occurs only when a specific histological variety grows in a syngeneic host to it. Furthermore, different types of *Y*_*nm*_ and *µ*_*T*_ values is related to unperturbed cancer with different types of symmetries, which can be observed in four possible situations. First, several cancer types growing in the same host (e.g., Ehrlich, Sa-37 fibrosarcoma, and F3II malignant tumors growing in BALB/c/Cenp mice) [7, 8, 74, 88]. Second, a given histological variety of cancer growing in different host types (e.g., breast cancer growing in different women or men) [69]. Third, the same type of unperturbed cancer with varying degrees of asymmetry (symmetric or asymmetric) [14]. Similarly, the same type of cancer in the same type of host, but one immunodeficient and the other immunocompetent [123]. It should be noted that unperturbed cancer in the experiment corresponds to control groups (organisms not under the influence of an external agent) [14, 26, 73, 88, 123].

In the experiment, Φ_1_(r, *θ, φ*), *ρ*_*v*_(*t*), *σ*_12_ and Turing spatiotemporal patterns are not only influenced by *Y*_*nm*_, *µ*_*T*_ and *t*, but also by type of unperturbed cancer histological variety; the pathway for cancer formation in the organism; viability of cancer cells; host type (e.g., human, mice, rat); host immunocompetence degree (immunodeficient and immunocompetent); and endogenous (e.g., emotional/psychological stress) and exogenous (e.g., environmental conditions) factors, according to the knowledge accumulated in preclinical [7, 8, 14, 23, 73, 74] and clinical [69, 89] studies.

We propose that dynamic self-regulations in unperturbed cancer is possible by means of an activator-inhibitor global mechanism in Turing pattern, two opposing dynamic intrinsic mechanisms that work synchronously but on different timescales and depend on the degree of *Y*_*nm*_ asymmetry, *µ*_*T*_ value and the time *t* during entire unperturbed TGK. The inhibition mechanism (blue color in Turing pattern) mobilizes negative charges (*ρ*_*v*_(*t*) *<* 0, red color) in the central region (symmetrical cancer) or in several non-central regions (asymmetrical cancer) throughout unperturbed cancer volume. Nevertheless, activator mechanism (red color in Turing pattern) mobilizes positive charges (*ρ*_*v*_(*t*) *>* 0, blue color) in these regions to self-regulate the electronegativity and symmetry of unperturbed cancer, avoid high instabilities throughout its entire volume, ensure its maximum survival, and guarantee its success in interaction and competence with the host. Therefore, inhibition mechanism (first term of Eq (26)) is associated to process that accelerates unperturbed TGK, whereas activation mechanism (second term of Eq (26)) is related to processes that decelerates it.

Although activation and inhibition mechanisms are well defined in Turing spatiotemporal patterns during unperturbed cancer growth, effects both mechanisms are contained in their respective global kinetics parameters *α* and *β* (Eq (26)). The later may suggest that these global kinetics parameters *α* and *β* in Gompertz formulations need to be reinterpreted from of the following two aspects. First, parameters *α* and *β* are a consequence of activator-inhibitor global mechanism in Turing pattern, which in turn are connected with mechanic, electric and mechanical-electrical parameters, as shown in this study. Second, parameters *α* and *β* are related with fractal dimension and mechanical properties in unperturbed TGK, as previously reported in [7, 13, 28, 41, 70, 90, 111, 115, 124–126].

The Φ_1_(r, *θ, φ*), *ρ*_*v*_(*t*), *σ*_12_ and Turing spatiotemporal patterns, depending on degree of asymmetry of *Y*_*nm*_ and *µ*_*T*_ value, and above discussed may suggest that negative and positive intratumoral regions correspond to depolarized and polarized regions that change dynamically in space-time according to the degree of asymmetry of *Y*_*nm*_, respectively. These depolarized regions are formed by depolarized cells and related to active regions of unperturbed cancer, whereas polarized regions are composed by polarized cells and associated with passive regions. Depolarized cells have low transmembrane potential (fast-dividing cells) and polarized cells have high transmembrane potential (non-dividing cells) [124].

We pressume that passive regions of unperturbed cancer are not inactive, like the central or non-central endogenous necrotic regions, but remain quiescent, which suggest that all its cancer cells do not participate in unperturbed cancer duplication process. These positive passive regions remain in quiescent states unless unperturbed cancer is significantly damaged by endogenous cell death mechanisms (necrosis, apoptosis, exfoliation, and metastasis) [69, 89] or the action of an external cytotoxic agent (e.g., anticancer therapies). For this, unperturbed cancer selectively and appropriately self-reverses the polarity from positive to negative in one or more quiescent intratumoral regions and their cells reactive and multiply again, which is possible because Φ_1_(r, *θ, φ*) = 0 mV (indicator of absence of ionic and biochemical activities) at the micro-interfaces between positive and negative intratumoral regions.

The existence of active and quiescent regions may justify why most aggressive unperturbed cancers, such as sarcomas, only duplicate 20 % of their cells and 80 % are in a state of dormancy [89]. The quantity and intensity of these active and dormant regions depend on the degree of asymmetry of *Y*_*nm*_ and *µ*_*T*_ value. A greater number of both negative and positive regions is observed for asymmetric odd *Y*_*nm*_, which correspond to unperturbed cancer more heterogeneous electrically and biologically and therefore more aggressive.

The above discussion suggest that intratumoral quiescent positive regions have a dual role depending on the needs of survival, growth, and metastasis of unperturbed cancer, as well as its protection against external cytotoxic agents. The first role could be associated with the self-regulation of symmetry, overall electronegativity, and all biophysical, chemical, and biological processes of unperturbed cancer, as mentioned above. The second role should be related to these dormant positive regions activating mechanisms that stimulate the growth and metastasis of unperturbed cancer in synergy with active negative regions.

Everything documented in this study may indicate that the anomalous bioelectricity of unperturbed cancer is not only related to metabolism, as [15, 127] report, but also has an essential role in the self-regulation of all its physical (e.g., electrical, mechanical, and thermodynamic), chemical (e.g., mass transfer), and biological (e.g., biological heterogeneity) processes, as well as in the complete unperturbed TGK to ensure self-symmetry, self-conservation of the global negativity of Φ_1_(r, *θ, φ*), *ρ*_*v*_(*t*) and *σ*_12_, self-regulations of negative (related to inhibitor mechanisms) and positive (linked to activator mechanisms) intratumoral regions, and self-maintenance of both electrical and biological heterogeneity of unperturbed cancer, as well as its maximum survival, growth, metastasis and protection against attack by external cytotoxic agents. Consequently, bioelectricity and biomechanics of unperturbed cancer should be given the greatest attention, as documented in other studies [13, 15–18, 20, 21, 23, 24, 37, 67, 68, 128–131], and its mechano-electrical nature should be considered as another of hallmarks of cancer [5], as suggested in [13].

## New insights and conclusions

Despite the results and hypotheses in this study, the anomalous bioelectrical connection of undisturbed cancer with its biology, entire TGK, and impacts on both different clinical variables in cancer organisms and parameters of anticancer therapies (oncospecific or under study) remain surprising, intriguing and fascinating even now. Therefore, further studies are required to delve deeper and understand better into these explicit relationships from primary changes in patterns of Φ_1_(r, *θ, φ*), *ρ*_*v*_(*t*), *σ*_12_ and Turing space-time patterns, and values of ||*ρ*_*v*_(*t*)|| and ||*ρ*_*vs*_(*t*)||, which depend markedly on the degree of asymmetry of *Y*_*nm*_ and *µ*_*T*_ value.

A better understanding of how *µ*_*T*_ and degree of *Y*_*nm*_ asymmetry are involved and growth and metastasis processes in unperturbed cancer, not addressed in this study, will be the subject of the next study of the serial results set (Part II). This will allow the connection between abnormal bioelectricity and biophysical-chemical in unperturbed cancer. Furthermore, the local and global space-time changes in the physical magnitudes Φ_1_(r, *θ, φ*), *ρ*_*v*_(*t*) and *σ*_12_ must be taken into account for the design and application of anticancer therapies, such as chemotherapy [69, 132], immunotherapy [113, 133] and physical therapies, essentially electrochemical ablation therapy (EA) [8, 14, 26, 52, 73, 88, 124] and irreversible electroporation (IRE) [132, 134]. These anticancer therapies may be applied to cancer separately or in combination (e.g., EA+immunotherapy, IRE+immunotherapy, EA+IRE, or EA+IRE+immunotherapy). This will be discussed in an another study of the serial results set (Part III).

Despite the connection between abnormal intratumoral bioelectricity and the overall behavior of unperturbed cancer, the results of this study have not addressed the connection between this bioelectricity and its components, such as the parenchyma (composed of neoplastic tumor cells) and the stroma (the tumor microenvironment consisting of tumor-induced supporting tissue, which includes blood vessels, immune cells, fibroblasts, and the extracellular matrix). This may be relevant for anticancer therapies, particularly those targeting specific molecular targets in cancer cells [135, 136] and/or constant and periodic external perturbations that act only on the host and not on the cancer, so that complete remission or long-term coexistence of host and cancer cell populations (cancer as a controlled chronic disease) is obtained [137]. The first group of anticancer therapies focuses on specific molecular targets crucial to cancer survival, such as: epidermal growth factor receptors (EGFR and HER2), protein kinases, vascular growth factors (VEGF), and immune checkpoints [135, 136, 138] [143]. This will be addressed in a future study of the serial results set (Part IV).

The understanding of unperturbed cancer anomalous bioelectricity should not only be approached from Φ_1_(r, *θ, φ*), *ρ*_*v*_(*t*), *σ*_12_ and Turing space-time patterns, and values of || *ρ*_*v*_(*t*) || and || *ρ*_*vs*_(*t*) || ; but also from biochemical-electrical processes at the cancer-surrounding healthy tissue interface, aspects that is related to the *ϵ* parameter, which could be connected to surgical margins (2 cm outside Σ) and therapeutic margins (2 cm outside the surgical margin) in clinics [69, 89]. The *ϵ* width increases when unperturbed cancer grows, as we corroborated theoretically in this study. Our model estimate 120 cells in this interface when *R*_*T*0_ = 5.6 mm; nevertheless, this model estimates 1000 and 2000 cells when unperturbed cancer surface is widened by 1 and 2 cm (*ϵ* width), respectively. Further studies are required to understand in depth the role of *ϵ* parameter in symmetry; overall electronegativity, specifically in the spherical cap and Σ; growth; angiogenesis; metastasis; and resistance against attack by external cytotoxic agents.

The results of this study suggest the need to experimentally verify the possible correspondence between intratumoral structural changes in space-times (e.g., active and necrotic intratumoral regions) and those of Φ_1_(r, *θ, φ*), *ρ*_*v*_(*t*) and *σ*_12_ spatiotemporal patterns in different solid cancer types, both in preclinical and clinical studies. This may be relevant to oncology because these intratumor structural changes in space-time are not tracked by pathological anatomy in a longitudinal study, since the animal/individual with cancer would require multiple biopsies or surgeries. This would lead to excessive manipulation of the same animal/individual, which would be counterproductive to bioethical considerations in humans and laboratory animals. Nevertheless, these intratumoral structural changes in space-time of both unperturbed and perturbed malignant tumors may be based on Φ_1_(r, *θ, φ*), *ρ*_*v*_(*t*) and *σ*_12_ spatiotemporal patterns, which can be measured experimentally with both magnetic resonance imaging and electric current density imaging, as suggest in [14].

The suggestions above in this section will allow for an understanding of the activation-inhibition process from both a biophysical and clinical perspective. It is unclear whether there is a single biological structure in unperturbed cancer that has a dual role, or whether different structures separately activate or inhibit its TGK. Furthermore, this study represents the groundwork for explicitly linking the bioelectricity of unperturbed cancer, by means of the magnitudes Φ_1_(r, *θ, φ*), *ρ*_*v*_(*t*) and *σ*_12_, not only with its global volume that grows over time, but also with other global parameters, such as its abnormal metabolism [16, 17] and global electrical impedance [23]. Furthermore, this bioelectricity could be linked to cancer biomarkers, whether biological [17, 18] or bioelectrical [23].

In conclusion, the anomalous bioelectricity and biomechanics in unperturbed cancer are due to intratumoral electrical heterogeneity and anisotropy according to Φ_1_(r, *θ, φ*), *ρ*_*v*_(*t*), *σ*_12_ and Turing spatiotemporal patterns, depending on the degree of *Y*_*nm*_ asymmetry, *µ*_*T*_ value 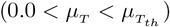 and *t*. Anomalous bioelectricity and biomechanics are closely related and both should be considered as another hallmark of cancer because these are essentially involved in self-regulation of symmetry, global electronegativity, intratumoral biological heterogeneity and anisotropy, growth, metastasis (by electrostatic and/or electromagnetic repulsion), abnormal metabolism of unperturbed cancer, as well as its protection from attack by cellular elements of the immune system and anticancer therapies (by formation of a heterogeneous electric shield), and in the appropriate and individualized selection of anticancer therapy, either alone or in combination.

## Acknowledgments

J.R.R.A, L.E.B.C. and R.A.B deeply appreciate the financial support of The Universidad Nacional Autónoma de México (UNAM) by the Dirección General de Asuntos del Personal Académico (DGAPA, Programa de Estancias de Investigación PREI, DGAP/DFA/2592/2025). L.E.B.C. thanks the Instituto de Investigaciones en Matemáticas Aplicadas y en Sistemas (IIMAS), UNAM, Mexico City, Mexico and the National Center for Applied Electromagnetism of the University of Oriente, Santiago de Cuba, Cuba, for their support and hospitality. J.R.R.A. thanks the Department of Mathematics and Mechanics of the IIMAS and DGAPA-PAPIIT UNAM under grant IN-109525. J.I.M.T. and L.E.B.C. gratefully acknowledge partial financial support for this study by MINECO, Spain, under the Project PID2022–141385NB–I00. J.B.R. gratefully acknowledges partial financial support for this study by Secretaría de investigación y posgrado del IPN, Project SIP20253412. The funders had no role in study design, data collection and analysis, decision to publish, or preparation of the manuscript.

## Supporting information

**S1 Video**. Biopotentials and volumetric charge density of unperturbed cancer for *Y*_2*m*_. Corresponding to Fig 4 of the main text.

**S2 Video**. Biopotentials and volumetric charge density of unperturbed cancer for *Y*_20_ for *µ*_*T*_ = 1.0. Corresponding to Figs 5A,B,C,D and 6A,B,C,D of the main text.

**S3 Video**. Biopotentials and volumetric charge density of unperturbed cancer for *Y*_21_ for *µ*_*T*_ = 1.0. Corresponding to Figs 5E,F,G,H and 6E,F,G,H of the main text.

**S4 Video**. Biopotentials and volumetric charge density of unperturbed cancer for *Y*_32_ for *µ*_*T*_ = 1.0. Corresponding to Figs 5I,J,K,L and 6I,J,K,L of the main text.

**S5 Video**. Biopotentials and volumetric charge density of unperturbed cancer for *Y*_20_ for *µ*_*T*_ = 25.0. Corresponding to Figs 7E,F,G,H and 8E,F,G,H of the main text.

**S6 Video**. Biopotentials and volumetric charge density of unperturbed cancer for *Y*_21_ for *µ*_*T*_ = 25.0. Corresponding to Figs 9A,B,C,D and 10A,B,C,D of the main text.

**S7 Video**. Biopotentials and volumetric charge density of unperturbed cancer for *Y*_32_ for *µ*_*T*_ = 25.0. Corresponding to Figs 9I,J,K,L and 10I,J,K,L of the main text.

**S8 Video**. Biopotentials and volumetric charge density of unperturbed cancer for *Y*_41_. Corresponding to Fig 11 of the main text.

**S9 Video**. Biopotentials and volumetric charge density for stripe patterns. Corresponding to Fig 14 of the main text.

## Author Contributions

**Conceptualization:** J. Roberto Romero-Arias, Luis Enrique Bergues Cabrales, José Alejandro Heredia Kindelán, Daniel Romero Rosales, Rafael A. Barrio, Juan Bory Reyes and Juan Ignacio Montijano Torcal.

**Formal analysis:** J. Roberto Romero-Arias, Luis Enrique Bergues Cabrales and Juan Ignacio Montijano Torcal.

**Funding acquisition:** Juan Ignacio Montijano Torcal.

**Investigation:** J. Roberto Romero-Arias, Luis Enrique Bergues Cabrales, José Alejandro Heredia Kindelán, Daniel Romero Rosales, Rafael A. Barrio, Juan Bory Reyes and Juan Ignacio Montijano Torcal

**Methodology:** J. Roberto Romero-Arias and Luis Enrique Bergues Cabrales.

**Software:** J. Roberto Romero-Arias.

**Visualization:** J. Roberto Romero-Arias, Luis Enrique Bergues Cabrales and Juan Ignacio Montijano Torcal.

**Writing – original draft:** J. Roberto Romero-Arias, Luis Enrique Bergues Cabrales and Juan Ignacio Montijano Torcal.

**Writing – review & editing:** J. Roberto Romero-Arias, Luis Enrique Bergues Cabrales, José Alejandro Heredia Kindelán, Daniel Romero Rosales, Rafael A. Barrio, Juan Bory Reyes and Juan Ignacio Montijano Torcal.

## Notes

### Competing Interest Statement

The authors have declared no competing interest.

